# Large-scale phenotyping and multi-trait selection identify low-methane Napier grass (*Cenchrus purpureus*) accessions under contrasting environmental conditions

**DOI:** 10.64898/2026.09.22.753521

**Authors:** Agalu W. Zeleke, Juan Palma-Hidalgo, Zaira Pardo Dominguez, Meki Muktar, Abel Teshome, Bayissa Hatew, Chris S. Jones, C. Jamie Newbold

## Abstract

Napier grass (*Cenchrus purpureus, NG*) is an important forage in tropical and subtropical livestock systems, but the extent of naturally occurring variation in its methane (CH_4_) production and its potential for selecting low-CH_4_ emitting genotypes remains poorly characterised. This study evaluated variation in CH_4_ yield among NG accessions, examined the effects of season and water stress, validated contrasting CH_4_ phenotypes across donor cows, and identified elite accessions combining low CH_4_ yield with favourable digestibility. A total of 750 NG samples representing 84 accessions grown under dry and wet seasons and moderate and severe water-stress were evaluated using a high-throughput *in-vitro* fermentation system. To further identify forages combining low-CH_4_ production with acceptable digestibility, a multi-trait selection approach was applied using biologically defined thresholds for CH_4_ yield and *in vitro* organic matter digestibility (IVOMD), followed by a weighted index assigning 60% weight to CH_4_ yield and 40% to IVOMD.

Methane yield varied widely among accessions, ranging from 0.02 to 5.07 mL/g DM and was affected by growing conditions. Methane production was generally lower during the wet season, while water stress was associated with changes in forage quality and fermentation characteristics. Low-CH_4_ producing accessions generally had lower IVOMD, crude protein and metabolizable energy and relatively higher fibre than high-CH_4_ producing accessions. Accessions initially classified as low and high-CH_4_ emitters based solely on CH_4_ yield were subsequently validated using rumen inoculum from three individual donor cows. Although the magnitude of CH_4_ production varied among donors, the contrast between low-and-high-CH_4_ accessions was generally retained, supporting the robustness of accession differences in CH_4_ production. Several accessions also maintained relatively low CH_4_ production across contrasting season and water-stress conditions, indicating that favourable CH_4_ phenotypes were not entirely environment dependent.

A multi-trait selection identified ten elite accessions that produced 0.019–0.810 mL CH_4_/g DM and had 55.2–69.0% IVOMD, whereas ten poor-performing accessions produced 2.80–3.89 mL CH_4_/g DM and had 50.1–54.7% IVOMD. Differences in hexose fermentation and CH_4_ produced per unit of fermented substrate further indicated contrasting fermentation efficiency between the groups. These results demonstrate substantial variation in NG and identify accessions with low-CH_4_ production, favourable digestibility and relatively stable performance across growing conditions. In conclusion, incorporating CH_4_ yield and digestibility into forage selection could support the development of lower-emission forage cultivars for sustainable climate-smart livestock production.

## 1. Introduction

Livestock, particularly ruminants, are one of the largest producers of greenhouse gases (GHG) worldwide. Methane (CH_4_) emissions from enteric fermentation and manure management by ruminant livestock are estimated to contribute to >32% of total anthropogenic GHG emissions globally (CCAC, 2021). Methane emissions are also linked to the loss of feed energy to animals, which was estimated at 2–12% of gross energy intake (Johnson and Johnson, 1995). Global anthropogenic CH_4_ emissions are expected to increase by more than 15% relative to 2010 levels by 2050, reaching almost 380 million tonnes per year (Höglund-Isaksson et al., 2020). To meet the increasing demand for animal-derived products while reducing anthropogenic GHG emissions from livestock, there is growing interest in the development of low-cost, sustainable and scalable forage-based mitigation strategies capable of reducing CH_4_ emissions, particularly in low-and-middle income countries without compromising animal productivity.

Napier grass (*Cenchrus purpureus,* NG) is one of the major tropical forage grasses, which grows widely across different agroecological zones in sub-Saharan Africa (Islam et al., 2023). Its ability to produce high biomass yield, perennial growth habit, fast regrowth after harvesting and tolerance to frequent cuttings have made NG one of the most important forage grass species for smallholder dairy production systems in the region (Habte et al., 2022; Muktar et al., 2022). In many tropical and subtropical countries, NG may comprise as much as 80% of cattle diets (Kabirizi et al., 2015), leading farmers to favour varieties with high biomass yields (40–59%, often linked to plant height) and rapid regrowth rates (typically 10–26%). However, in Ethiopia, there is large variation in agronomic performance (e.g., dry matter yield, leaf-to-stem ratio) among NG accessions across diverse agroecological environments. While information regarding agronomic traits and yield performance stability of NG accessions has been reported from different agro-ecologies in Ethiopia (Adnew and Asmare, 2023; Kebede et al., 2017; Mossie et al., 2024), little is known about the variation in CH_4_ production from genetically diverse NG accessions across environments. Consequently, characterising diverse NG accessions for their CH_4_ emission potential (*in vitro*) and associated ruminal traits is essential to support the development of phenotypic parameters that could enable the breeding of low-CH_4_-emitting forages.

Recent advances in high-throughput *in vitro* fermentation techniques enable rapid and cost-effective evaluation of hundreds of forage samples under controlled laboratory conditions while substantially reducing labour, incubation space, reagent use, and analytical time (Mauricio et al., 1999; Pellikaan et al., 2011). With these fermentation systems, the substrates and buffered rumen fluid are directly incubated within sealed 20 mL GC vials (leaving headspace for gas accumulation), and the headspace CH_4_ concentration can be determined immediately following incubation, thereby improving experimental throughput and handling efficiency. These systems are thus becoming more important for large-scale forage phenotyping and for identifying feed resources with low CH_4_-emitting traits, contributing to climate-smart livestock production systems. The aims of this study were 1) to evaluate variation in CH_4_ production among a large collection of NG accessions from International Livestock Research Institute (ILRI) forage gene bank, 2) to assess the effects of season and water stress on CH_4_ production and rumen fermentation characteristics, and 3) to validate the consistency of CH_4_ phenotypes across individual donor animals. Specifically, to identify accessions combining low CH_4_ emissions with adequate digestibility, which could be used by breeders, and farmers to mitigate the environmental footprint of ruminant production in smallholder systems in low-income countries.

## 2. Materials and methods

### 2.1. Plant materials and growing conditions

Eighty-four (n = 84) NG accessions were sourced from ILRI forage gene-bank and cultivated under field conditions at the Bishoftu research site, Ethiopia (008°47′20″ N and 038°59′15″ E), from 2018 to 2020 for two seasons (Wet [June to September] and Dry [November to May]) in four crop blocks. The altitude of Bishoftu is 1890 m.a.s.l., with an Alfisol soil type. The origin and diversity of these genotypes were previously described by Muktar et al. (2019). Within each growing season, plants were subjected to two water regimes: moderate water stress (MWS), maintained at 20% soil volumetric water content (VWC), and severe water stress (SWS), maintained at 10% soil VWC. The soil moisture content of the field plots was monitored using a Delta soil moisture probe (HD, UK), and details of field establishment, land preparation, and the physical and chemical properties of the experimental soil have been described previously by Habte et al. (2022). At 3-months after planting, all plants were cut to a height of 50 mm above ground level. Subsequently, harvesting and data collection were conducted following every 8 weeks of regrowth.

A total of 750 forage samples comprising whole plants (leaf and stem) were collected across seasons and water-stress treatments for further analysis. Following collections, samples were oven-dried at 60°C for 48 h, ground to pass through a 1-mm sieve, and sealed in labelled polythene grip-seal bags bearing unique sample identifiers. Detailed information on accession origins (metadata), experimental design, field management practices, growing conditions, data collection and measurements have been previously reported by Muktar et al. (2019). Subsequently, approximately 5–10 g of each dried sample was shipped to Scotland’s Rural College (SRUC, United Kingdom) for high-throughput *in vitro* incubation and CH_4_ analysis. Samples were stored at room temperature (<25°C) until analysis. For CH_4_ quantification, 45 mg of each accession was weighed into a 20 mL Agilent clear headspace vial. Subsamples of the dried material were analysed for feed quality traits, including neutral detergent fibre (NDF), acid detergent fibre (ADF), acid detergent lignin (ADL), organic matter (OM), crude protein (CP), *in vitro* organic matter digestibility (IVOMD) and metabolisable energy (ME) as reported by Habte et al. (2020), with these traits expressed on a percentage of dry matter (DM) basis, except for ME, which is expressed as MJ/kg DM.

### 2.2. High-throughput *in vitro* fermentation assay

#### 2.2.1. Buffered rumen fluid preparation

Rumen fluid (RF) samples were collected from three rumen-cannulated Jersey cows, aged 5–12 years, grazing a sward of perennial ryegrass *ad libitum* with a garlic lick to utilise at their leisure. Fresh rumen contents were transferred from each cow into separate pre-warmed insulated one-liter thermos flasks and transported to the lab within 1 h of collection. The contents from each cow were filtered through three layers of cheesecloth, pooled, and diluted with Menke and Steingass (1988) buffer solution in a 1:2 ratio (buffer:RF) as described by Khan and Chaudhry (2021). With the buffer continuously flushed with CO2, the desired RF quantity was added as soon as the pink color disappeared. The buffer solution consisted of NaHCO_3_ (35 g) and (NH_4_) HCO_3_ (4 g) dissolved in 1-L distilled water. The macro mineral solution contained Na_2_HPO_4_ (5.7 g), KH_2_PO4 (6.2 g) and MgSO_4_*7H_2_O (0.6 g). An additional 120 µL micromineral solution and 1.22 mL resazurin/L distilled water were used. The reducing solution contained sodium sulfide nonahydrate (Na₂S·9H₂O; 0.313 g/L) and was added before incubation to maintain anaerobic conditions in the vessel.

#### 2.2.2. In vitro incubation

Napier grass accessions were evaluated using a 24 h *in vitro* batch culture system designed to mimic the rumen. Samples were randomly allocated into four incubation batches, each representing all four crop blocks and both growing seasons. The buffered inoculum was maintained at 39°C under continuous CO₂ flushing to preserve anaerobic conditions. Aliquots (5 mL) of the buffered RF were anaerobically dispensed into 20 mL incubation vials containing 45 mg of ground NG sample. With the vials flushed by CO₂ gas, they were equipped with butyl rubber stoppers and aluminum crimps, gently agitated, and then incubated at 39°C for 24 h. Throughout the incubation setup, buffers, the rumen fluid mixture, and the vials were continuously flushed with CO₂ gas, thereby ensuring anaerobic conditions. Blank vials containing only buffered RF were included in each run as controls, and one sample per batch was included at the start and end of each incubation run to verify consistency and quality control.

#### 2.2.3. Methane quantification

After 24 h of incubation, vials were removed from the incubator, and total gas production was recorded using a pressure transducer (Vernier LabQuest 2, LQ2-LE). Fermentation was then terminated by adding 1.1 mL of 20% orthophosphoric acid containing 2-ethylbutyric acid as an internal standard for subsequent analyses of CH_4_ and volatile fatty acids (VFA). Methane concentration was determined immediately using a gas chromatograph (Agilent 8890 GC System) equipped with an automated headspace sampler (Agilent 8697). Separation was achieved on an HP-5 capillary column (30 m length × 0.320 mm internal diameter × 0.25 µm film thickness, Agilent Technologies USA) using nitrogen as a carrier gas. Following CH_4_ measurement, 1.6 mL of fermentation medium from each vial was transferred into microcentrifuge tubes and stored at −20°C for subsequent VFA analysis. A standard curve for CH_4_ quantification was generated using four calibration CH_4_ concentrations (2.45%, 5.13%, 10.25%, and 15.08%), each prepared in 20 mL GC vials and analysed in quintuplicate (n = 5), resulting in 20 standards per run. Individual CH_4_ concentrations were then calculated using the slope and intercept derived from the calibration curve.

#### 2.2.4. Volatile fatty acid (VFA) analysis

For VFA analysis, samples were thawed at room temperature and centrifuged at 10,000 × g, 4°C for 10 min. The resulting supernatant was filtered through nylon syringe filters (0.2 µm pore size, 25 mm diameter, Fisher scientific) and transferred into 1.5 mL gas chromatography (GC) vials, which were then sealed and VFA concentrations were quantified using an Agilent 6850 GC system for on-column injection and flame ionisation detection in accordance with the manufacturer’s instructions. For separation, a HP-FFAP column 30 m × 0.53mm i.d × 1 µm film thickness was employed. Calibration curves were generated using Supelco_®_ volatile free acid mix (certified reference material Sigmaaldrich, CRM46975, USA), which included acetic, propionic, iso-butyric, butyric, iso-valeric and valeric acids. Four concentrations were used (2, 4, 6, and 8 mM): 2 mM = 200 µL VFA mix + 200 µL internal standard + 600 µL distilled water, 4 mM = 400 µL VFA mix + 200 µL internal standard + 400 µL distilled water, 6 mM = 600 µL VFA mix + 200 µL internal standard + 200 µL distilled water and 8 mM = 800 µL VFA mix + 200 µL internal standard). Concentrations of VFA were calculated from corrected chromatographic peak areas (peak area ratio of compound of interest to internal standard) using the slope and intercept of the calibration curve. Final VFA concentrations were subsequently corrected for dilution (factor = 0.82) to account for the addition of the internal standard.

### 2.3. Estimation of hexose fermentation and methane production intensity

The amount of carbohydrate fermented was estimated from the major end-products of fermentation (VFAs) using a stoichiometric approach based on rumen fermentation pathways. The concentration of estimated hexose fermented (HF) was calculated from the molar concentrations of major VFAs according to the stoichiometry approaches described by Wolin (1960) and Della Rosa et al. (2026) using the equation below:

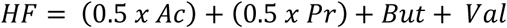

*Where HF = hexose fermented, Ac =acetate, Pr = propionate, But = butyrate and Val = valerate*.

The calculated HF represented hexose fermentation equivalents in the incubation medium (mM). Total HF was obtained by multiplying HF concentration by the incubation liquid volume (0.005L). Hexose fermented per gram of incubated substrate was calculated by dividing total HF by the amount of incubated substrate (0.045 g). Methane intensity was then calculated as the amount of CH_4_ produced per unit of estimated HF, calculated as the ratio of CH_4_ yield (mmol/g DM) to estimated HF (mmol/g DM) derived from major VFA end-products.

### 2.4. Validation of extreme CH_4_ phenotypes across animals

To evaluate the reproducibility of CH_4_ production across animals, 10 NG accessions with the highest CH_4_ emissions and 10 with the lowest CH_4_ emissions were selected (as described below). The RF was collected separately from three rumen-cannulated cows and treated as independent biological replicates. The validation assay followed the same *in vitro* fermentation protocol described above, with minor modifications. Briefly, 300 mg of each NG accession was weighed into 120-mL Wheaton serum bottles and inoculated with 30 mL of buffered RF (20 mL buffer and 10 mL RF), followed by incubation at 39°C for 24 h.

After incubation, total gas production was measured using a pressure transducer as previously described. Gas samples were collected using a Terumo™ Agani™ 18 G × 1.5″ needle attached to a 20 mL polypropylene Luer-lock syringe and transferred into pre-vacuumed 20 mL Agilent clear headspace crimp vials, which were immediately sealed with PTFE/silicone septa aluminium caps. Methane concentration was subsequently quantified using an Agilent 8890 GC system as described above. Following gas sampling, 1.6 mL of fermentation fluid was aliquoted into a 2-mL microcentrifuge tube and stored at −20°C for subsequent VFA analysis. Samples for VFA analysis were thawed at room temperature, centrifuged at 10,000 × g, 4°C for 10 min, and analysed using Agilent 6850 GC system following the manufacturer’s instructions as described above.

### 2.5. Statistical analysis

#### 2.5.1. Screening model

All data, except data from the validation test, were analysed in R (version 4.5.1, RStudio version 2026.01.0) using a linear mixed model using the ‘lmer’ function from ‘lmerTest’ package in R. A full 3-way interaction structure (Season × Stress × Accession) was initially fitted in this model but was later removed because it did not improve model fit and most parameters were not statistically significant (p > 0.05). The final model was fitted with a 2-way interaction (as presented below), with season, water stress and accession as the main fixed effects, block (replicate) as a random variable and overall error as sources of variation using the following equation:

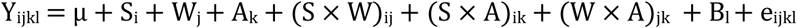

*Where Y_ij_ = response variable, μ = overall mean, S_i_ = fixed effect of season, W_j_ = fixed effect of water stress, A_k_ = fixed effect of accession, (S × W)_ij_ = interaction effect of season and water stress, (S × A)_ik_ = interaction effect of season and accession, (W × A)_jk_ = interaction effect of water stress and accession, and B_l_ = block fitted as a random effect, and e_ijkl_ = residual error.*

After fitting a linear mixed model, estimated marginal means (EMMs) of CH_4_ yield for each accession were calculated using the ‘emmeans’ function in R. Accession means were calculated by averaging predicted responses across each season-water stress combination while retaining the accession-specific interaction effects included in the model, thereby providing adjusted accession means that account for environmental variation. All accessions were ranked from highest to lowest based exclusively on CH_4_ yield (mL/g DM) and the top 10 (high CH_4_-yielding) and bottom 10 (low CH_4_-yielding) accessions were selected for further analysis of differences in fermentation and nutritional-related characteristics among the two groups. The non-parametric Wilcoxon rank-sum test was employed to analyse differences in CH_4_ yield (mL/g DM), gas volume (mL), CP, NDF, IVOMD, acetic, propionic, butyric, valeric and acetic: propionic between extreme phenotypes (top 10 and bottom 10 accessions), due to violations of normality and homogeneity of variance assumptions, with CH_4_ group (high vs low) as the main fixed effect:

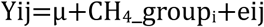

Where Y_ij_ = response variable, μ = overall mean, CH4group_i_ = fixed effect of CH4-emitting potential of accessions (high emitters vs low emitters) and e_ij_ = residual error.

#### 2.5.2. Validation model

The top 10 (high CH_4_-yielding) and bottom 10 (low CH_4_-yielding) accessions as described above were further validated across three individual cows and data from this validation trial were analysed using a mixed model with accession as the primary fixed effect and animal as a random effect:

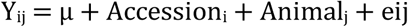

*Where Y_ij_ = response variable, μ = overall mean, Accession_i_ = fixed effect of accession, Animal_j_ was fitted as a random effect, eij = residual error.*

Subsequently, the relationship between CH_4_ yield measured in the screening trial (screening CH_4_ yield) and that measured in the validation trial (validation CH_4_ yield) was assessed in both high-and low-CH_4_ yielding accessions, with this association visualised using a linear regression plot created in R with the ‘ggplot2’ package (Wickham, 2016).

To further select accessions with extreme phenotypes that possess both relatively low CH_4_ and acceptable IVOMD, a multi-trait selection index was performed. First, all accessions were categorised based on biologically meaningful thresholds as elite (CH_4_ ≤ 1.5 mL/g DM and IVOMD > 55%), poor (CH_4_ > 1.5 mL/g DM and IVOMD ≤ 55%) and intermediate (accessions that were neither elite nor poor). Second, a weighted, standardised selection index (60% CH_4_ + 40% IVOMD) was calculated for accessions in the elite and poor groups to identify genotypes that mitigate CH_4_ emissions while maintaining adequate digestibility. The higher weight assigned to CH_4_ yield reflects its greater importance, as mitigating CH_4_ emission was the primary aim of the selection, while IVOMD was included to avoid choosing low-CH_4_-emitting accessions with poor nutritional value. Subsequently, the top 10 most promising accessions (high IVOMD and low-CH_4_ yield, Elite) and the bottom 10 least desirable accessions (low IVOMD and high-CH_4_ yield, Poor) were selected for further discussion (Supplementary Fig. 1). An IVOMD threshold of > 55% was used to identify accessions with moderate-to high-digestibility, as digestibility values above 55% have been considered as appropriate for good quality tropical forage with increased concentrations of fermentable organic matter (Van Soest, 1994; Minson, 2012). Similarly, a CH_4_ yield threshold of ≤ 1.5 mL/g DM was used to identify low-CH_4_-producing accessions while still maintaining an acceptable level of digestibility, with this threshold representing accessions in the lower quartile of our observed CH_4_ distribution. Significant differences between the two groups were determined using the Wilcoxon rank-sum test as described above.

To explore whether CH_4_-emission phenotypes were consistent across growing conditions, accessions were assigned to CH_4_-emission categories for each treatment combination (Dry-MWS, Dry-SWS, Wet-MWS, Wet-SWS). Within each treatment, CH_4_ yield values were ranked and divided into three groups of equal size (tertiles) using the ‘ntile’ function in R. Accessions in the first third of the CH_4_ yield distribution were assigned as ‘low’ CH_4_ emitters, middle third accessions were ‘medium’ emitters, and accessions in the top third were ‘high’ emitters. One accession that did not have values for all four growing conditions was excluded from further analysis. An alluvial plot created using the ‘ggalluvial’ package in R was used to visualise transitions in CH_4_-emission classification between growing conditions.

Pearson’s correlation coefficients and significance between fermentation parameters and feed quality variables was determined using the ‘cor.test’ function in R and visualised as a heatmap. Principal component analysis (PCA) was performed based on Euclidean distance matrices using the vegan package in R to assess clustering patterns of NG accessions across season, growing condition and extreme CH_4_ phenotypes. Significant differences were declared at *p* < 0.05; trends were defined as 0.05 < *p* < 0.10.

## 3. Results

### 3.1. Variation in methane production among NG accessions

Methane yield across accessions and growing conditions ranged from 0.02 to 5.07 mL/g DM (Supplementary Table 1), while the mean CH_4_ yield was 1.92 ± 0.90 mL/g DM. Within the dry season CH_4_ yield varied from 0.03 (under MWS) to 5.07 (SWS) mL/g DM, with the mean of 2.20 ± 0.70 and 2.41 ± 0.89 mL/g DM under MWS and SWS conditions, respectively. Wet season CH_4_ yield was generally lower than dry season irrespective of water stress levels. The CH_4_ yield during this season ranged from 0.02 to 3.89 mL/g DM and averaged at 1.56 ± 0.84 mL/g DM across both MWS and SWS conditions. Methane yield showed a considerable variation among NG accessions, with an overall coefficient of variation (CV) of 47%.

### 3.2. Effects of season, water stress, accession and their interaction on methane and VFA production

Results from a linear mixed-effects model showed that season, accessions and season × water stress interaction had significant effects on CH_4_ yield (p < 0.01, Fig. 1), while water stress (p = 0.185), water stress × accessions (p = 0.969), and season × accessions (p = 0.375) interactions were not significant. Block (replicate) as a random effect showed moderate variance (0.291), with moderate unexplained variability among experimental units (residual error = 0.427). We observed that CH_4_ production was typically higher during dry periods than the wet season, with the SWS in the wet season resulting in the greatest reduction (Fig. 2A).

**Fig. 1.**
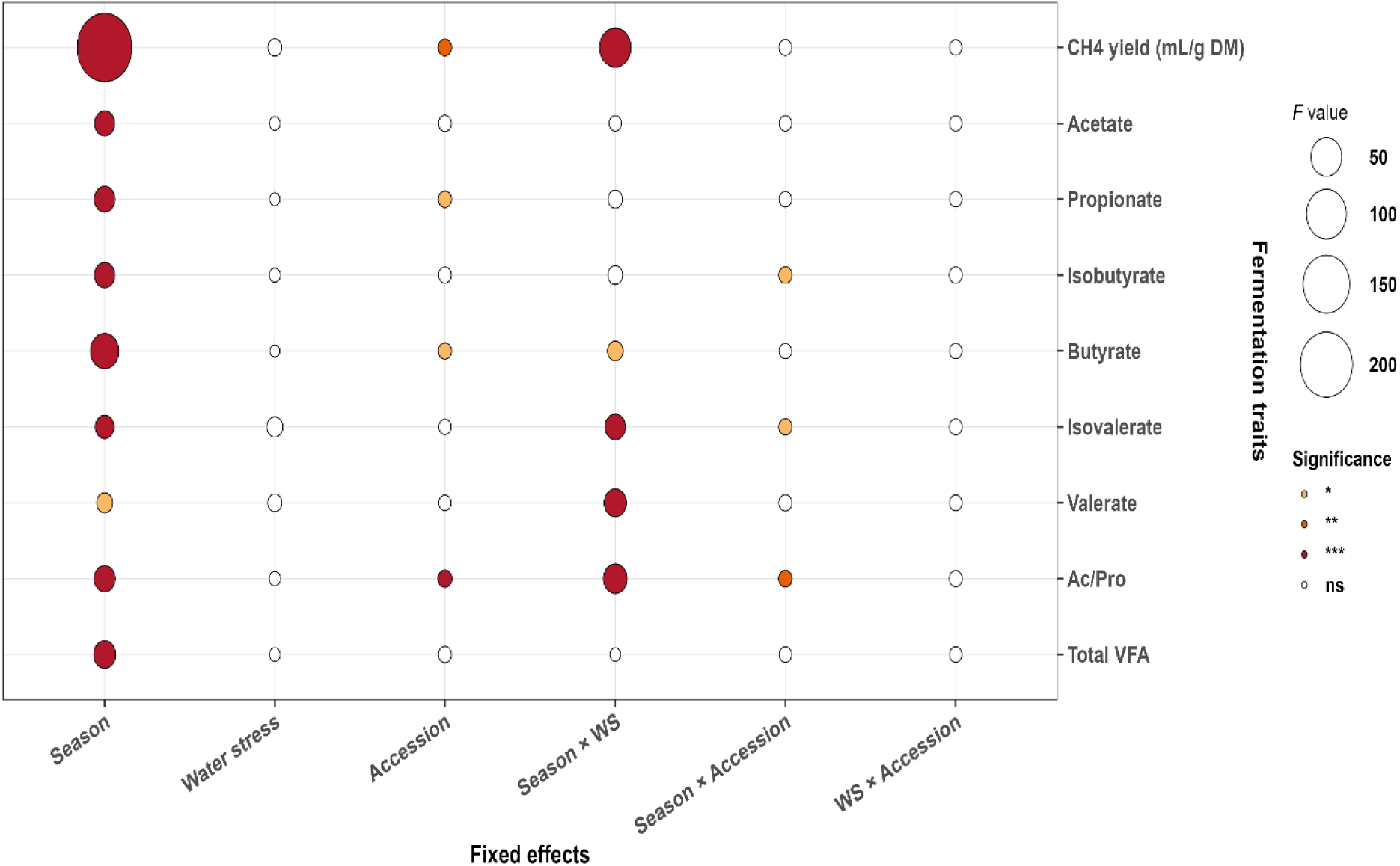
Effects of season, water stress, and accession and their interactions on fermentation related traits based on Type III ANOVA (linear mixed-effects models). Fixed effects: season, water stress, accession, season × WS, season × accession, and WS × accession. Abbreviations: WS = water stress, CH_4_ = methane yield (mL/g DM), VFA = volatile fatty acid, Ac/Pro = acetic to propionic ratio. Significance levels: ns = non-significant, * p < 0.05, ** p < 0.01, *** p < 0.001.

**Fig. 2.**
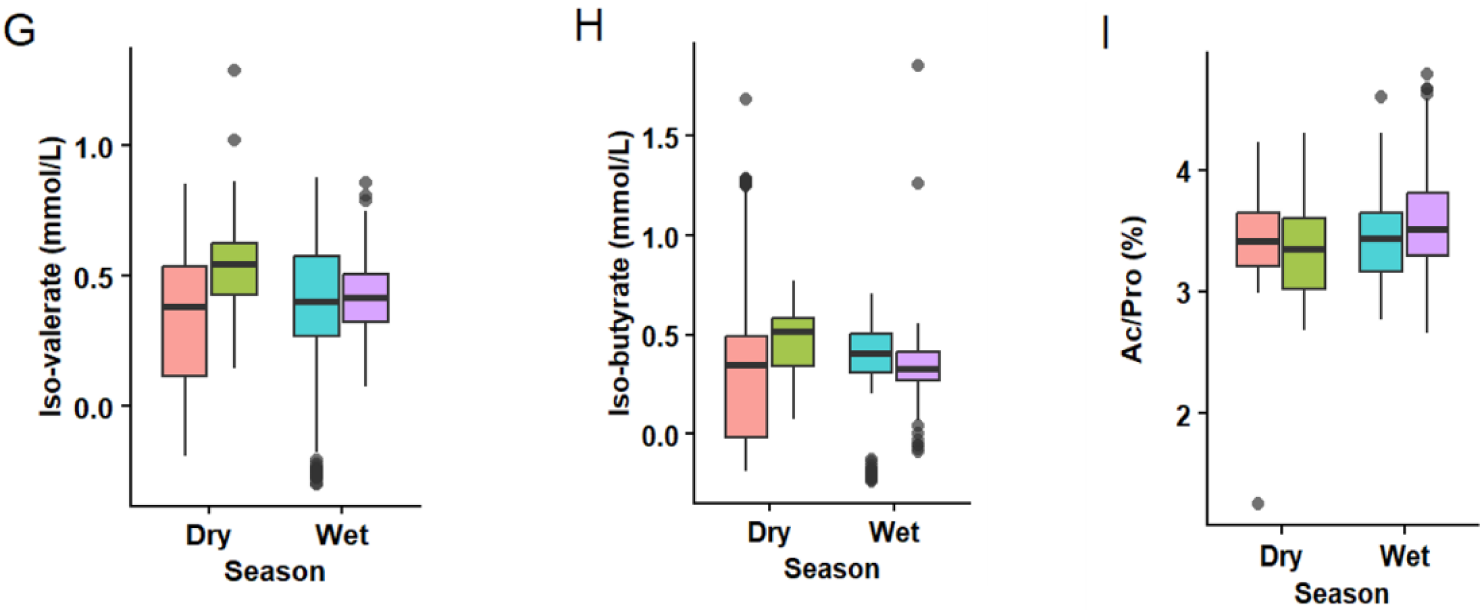
Boxplot showing distribution and variability of CH_4_ yield (A), total VFA (B), acetate (C), propionate (D), butyrate (E), valerate (F), iso-valerate (G), iso-butyrate (H) concentrations and acetate: propionate (I) across seasons and water stress conditions. MWS = moderate water stress, SWS = severe water stress.

While season × accessions interaction was not significant, some accessions responded differently between seasons (Fig. 1). For example, accessions 14389, 16796, 16813, 16835, BAGCE 81 and CNPGL 00-1-1 decreased CH_4_ by 1.49, 1.40, 1.63, 1.35, 2.05 and 2.01 mL/g DM, respectively, during wet season compared to dry season (p *<* 0.05). The significant negative interaction coefficient for season × water stress showed that CH_4_ yield was reduced by 0.69 mL/g DM during the wet season under SWS condition compared to the dry season (p < 0.001).

EMMs further demonstrated variability among accessions across season and water-stress treatments, with accessions 16819, CNPGL 00-1-1, 16804, 16813, and 16836 exhibiting the highest EMMs of 3.34, 3.34, 3.31, 3.26 and 3.25 mL/g DM, respectively, while accessions BAGCE 93, 18438, and 18448 had the lowest EMMs of 0.95,1.60, and 1.65 mL/g DM, respectively, during the dry season under MWS (Dry–MWS). When evaluating results from the wet season with MWS (Wet–MWS), the overall estimates decreased from those recorded under Dry–MWS; however, some accessions still recorded relatively high estimates, including BAGCE 53, 16806, 16786, 16804, and 16836. Four accessions 16788, 16791, BAGCE 81 and 16621, had the lowest EMMs of 1.21, 1.32, 1.41 and 1.45 mL/g DM, respectively, under Wet–MWS.

Under dry season with SWS (Dry–SWS), accessions CNPGL 92-133-3, BAGCE 86, 15357, 16819, and 16796 had the highest EMMs of 3.46, 3.15, 3.11, 3.03 and 2.98 mL/g DM, respectively, whereas accessions BAGCE 93, 18448, and 16782 showed relatively low EMMs of 1.54, 1.58 and 1.58 mL/ g DM, respectively under Dry–SWS. Wet season combined with SWS (Wet–SWS) had the lowest grand mean EMMs across all accessions. Accessions that had the highest EMMs under this treatment included 16840, 16787, and CNPGL 92-133-3, with mean estimates of 2.04, 1.98, and 1.97 mL/g DM, respectively, while accessions CNPGL 00-1-1, BAGCE 81, 14982, 16621 and 16813 had the lowest EMMs. Generally, there were some accessions that had relatively high EMMs across water stress conditions, including CNPGL 92-133-3, BAGCE 86, 16819, 16796, and 15357 regardless of season. Accessions like BAGCE 93, 14982, 16621, and CNPGL 00-1-1 tended to be more affected by Wet–SWS. While total VFA concentration remained unchanged across water stress, accession, season × water stress, and season × accession interactions (p > 0.05, Fig. 1), seasonal-specific variations were observed (p < 0.001), with the mean values ranging from 50.5 to 54.2 mmol/L during wet and dry season, respectively (Fig. 2B). Similar trends were observed for acetate (Fig. 2C), propionate (Fig. 2D) and butyrate (Fig. 2E) concentrations, with corresponding mean values ranging from 34.9 to 36.7 mmol/L, from 10.3 to 11.1 mmol/L, and from 4.5 to 5.1 mmol/L during the wet and dry seasons, respectively (p < 0.001). Similarly, season was the dominant factor influencing the concentration of valerate (Fig. 2F), iso-valerate (Fig. 2G), and iso-butyrate (Fig. 2H), with significant season × water stress interactions for valerate, and iso-valerate, and season × accession for iso-butyrate concentrations (p < 0.05, Fig. 1). The concentrations of butyrate, valerate and isovalerate reduced by 0.39, 0.17, and 0.12 mmol/L, respectively, during the wet season under SWS conditions (p < 0.05). The ratio of acetate:propionate was also affected by season (p < 0.001, Fig. 2I), and interactions of season × water stress (p < 0.001), and season × accession (p < 0.01, Fig. 1), with the ratio decreased by 0.25 percentage units during the wet season under SWS conditions (p < 0.001).

Several accessions exhibited season-specific effects on VFA concentration (p < 0.001). For example, acetate decreased by 21.32, 23.14, 20.58, 22.41, and 23.49 mmol/L in accessions 16807, 16816, 16819, BAGCE 17, and CNPGL 93-01-1, respectively (p < 0.001, supplementary Table 2) during the wet season compared with the dry season. The same accessions showed consistently lower propionate concentration of 8.47, 7.47, 7.09, 7.90, and 7.69 mmol/L (supplementary Table 3) and lower butyrate concentrations of 3.30, 3.24, 3.21, 3.05, and 3.02 mmol/L, respectively, during the wet season compared with the dry season (p < 0.001, supplementary Table 4).

### 3.3. Relationships between methane production and fermentation characteristics

The relationship between CH_4_ production, VFA profiles, and feed quality traits was assessed using Pearson correlation and visualised as heatmaps (Fig. 3). The CH_4_ yield, total gas, and CH_4_ yield per fermented hexose were all negatively correlated with NDF (r = - 0.26,-0.26, and-0.23, respectively) and with ADF (r = - 0.29,-0.29 and-0.26, respectively) (p < 0.001). In contrast, these CH_4_-related traits were positively associated with IVOMD and ME (p < 0.001, Fig. 3). As expected, CH_4_ yield had a strong positive association with total gas production (r = 0.92, p < 0.001). Additionally, CH_4_ yield showed a moderate positive correlation with total VFA (r = 0.54, p < 0.001), acetate (r = 0.51, p < 0.001), propionate (r = 0.54, p < 0.001), butyrate (0.56, p < 0.001), and total hexose fermented (r = 0.54, p < 0.001). Interestingly, CH_4_ yield showed a moderate negative relationship with acetate-to-propionate ratio (Ac/Pro; r =-0.41, p < 0.001).

**Fig. 3.**
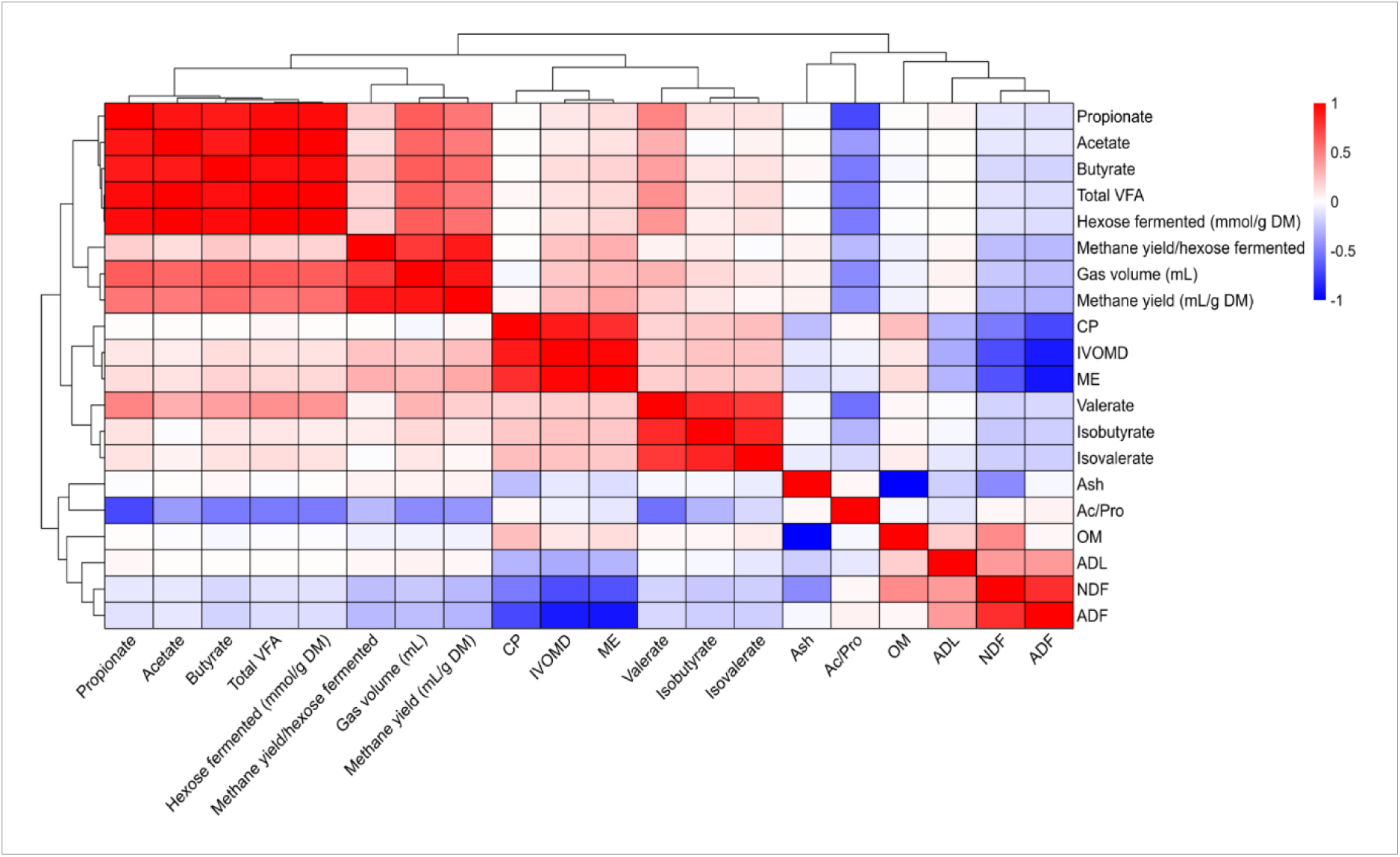
Correlation heatmap showing the relationships between CH_4_ production, VFA and feed quality related traits across NG accessions. ADF = acid detergent fibre, NDF = nutrient detergent fibre, ADL = acid detergent lignin, IVOMD = *in vitro* organic matter digestibility, ME = metabolisable energy, CP = crude protein, Ac/Pro = acetate to propionate ratio.

### 3.4. Identification of low-CH_4_ producing Napier grass accessions

The top 10 (high CH_4_ yielding) and bottom 10 (low CH_4_ yielding) accessions were selected to further examine differences in fermentation patterns and feed quality traits among these two extreme phenotypes. As shown in Table 1, there was wide variability in CH_4_ yield among NG accessions grown under contrasting seasons and water stress conditions. Most of the high CH_4_ yielding accessions were recorded during the dry season under SWS conditions (ranging from 3.89 to 5.07 mL/g DM), whereas low CH_4_ yielding accessions were mostly under wet season and SWS conditions (varying from 0.019 to 0.072 mL/g DM). Significant differences were observed between high-and low CH_4_-producing accessions for several feed composition and fermentation characteristics (Table 2). For example, most of the high CH_4_ yielding accessions had higher gas production (p < 0.001), IVOMD (p = 0.004), acetate (p < 0.001), propionate (p < 0.001) and butyrate concentrations (p < 0.001) than the low CH_4_ yielding accessions. Conversely, low CH_4_ producing accessions exhibited lower digestibility, fermentation activity and VFA production, potentially associated with higher NDF and ADF (p = 0.007 and p = 0.004, respectively). Contrary to expectations, acetate:propionate ratio was higher in high CH_4_ producing accessions compared with low CH_4_ producing accessions (p = 0.001).

**Table 1.**
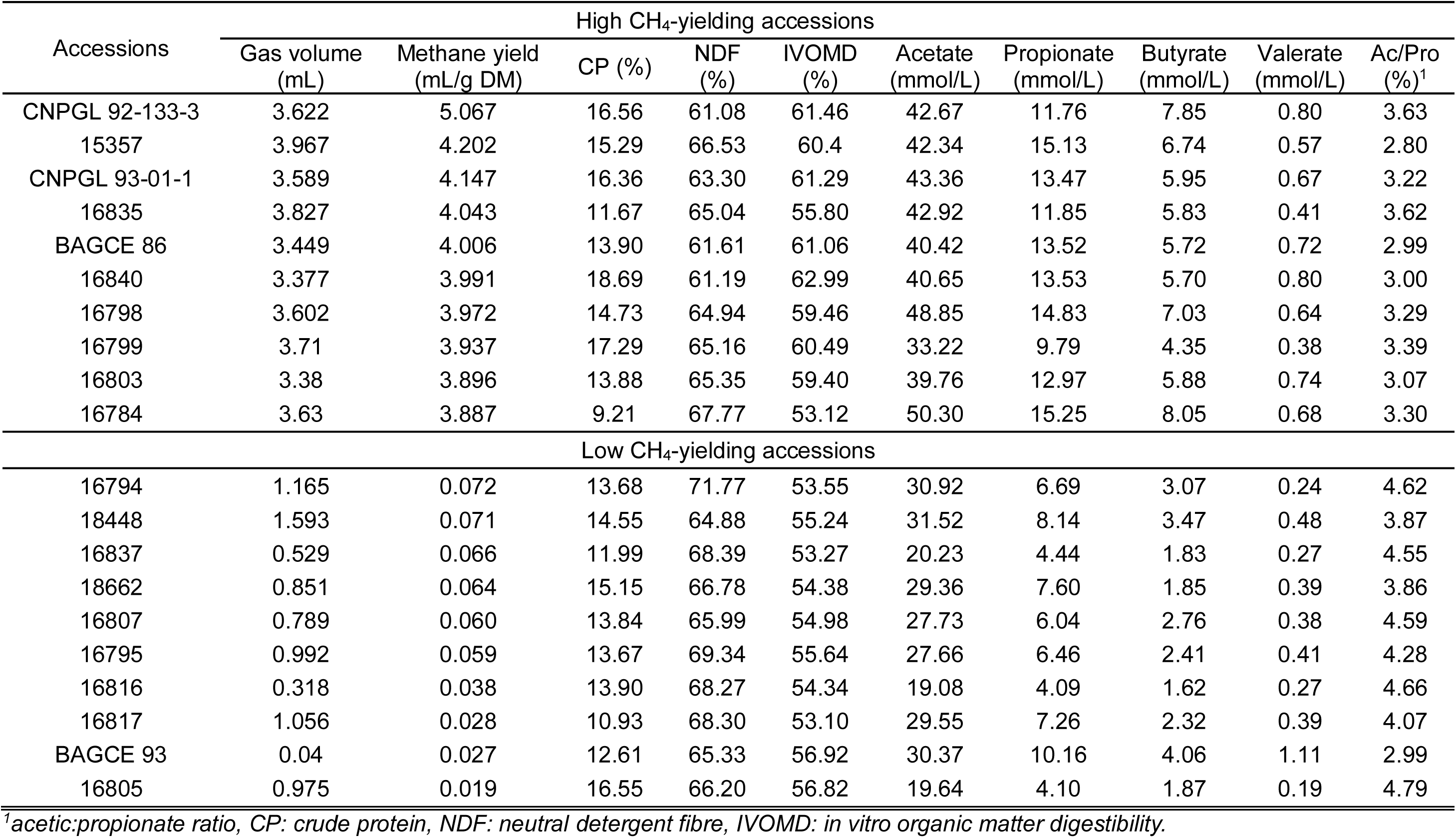
Top and bottom 10 NG accessions ranked by CH_4_ yield (mL /g DM) under contrasting seasonal and water stress conditions.

**Table 2.**
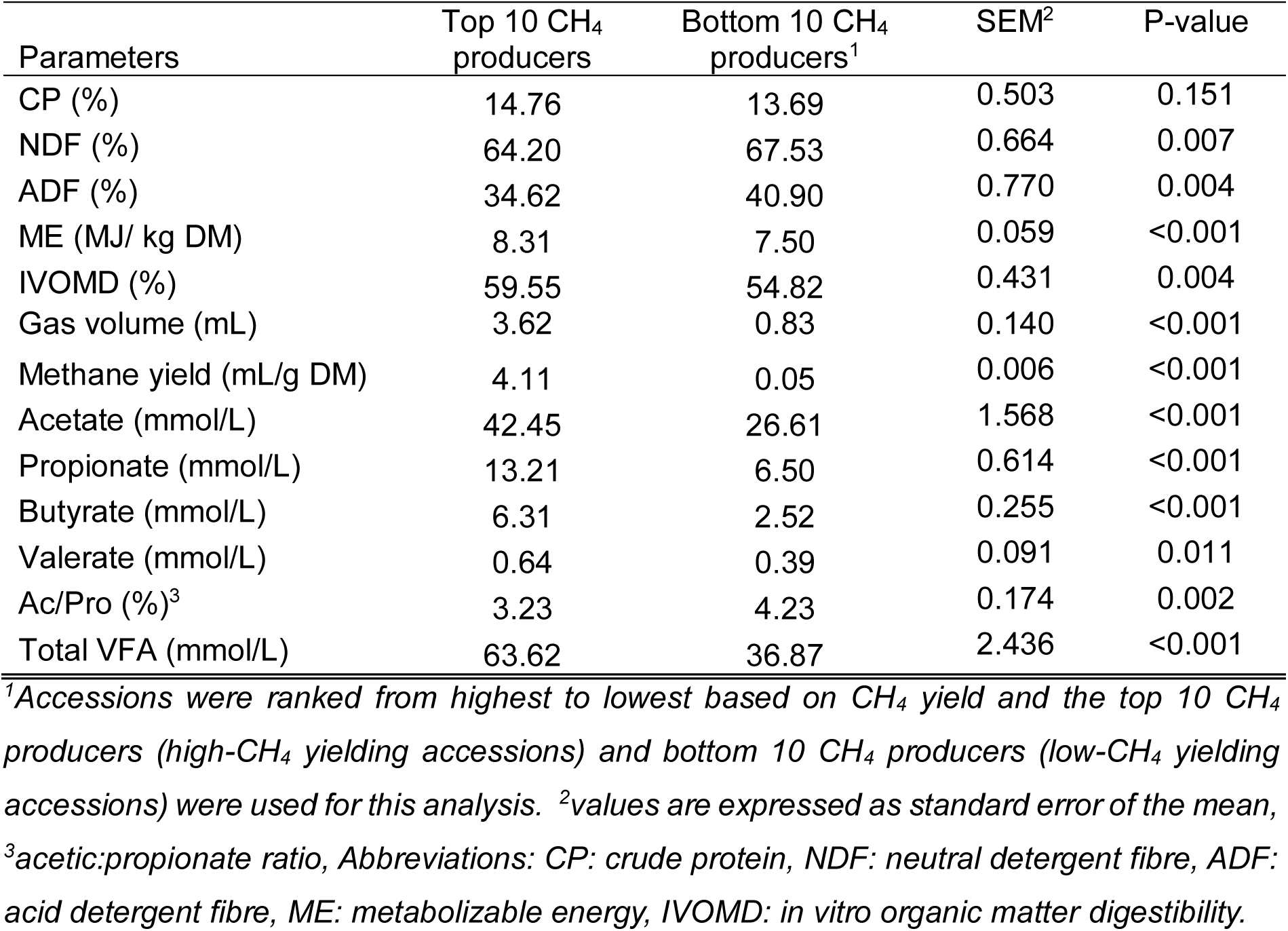
Mean compositional differences between top and bottom CH4 groups.

Principal component analysis was performed to evaluate the multivariate difference between high and low CH_4_-producing accessions based on CH_4_ production as well as fermentation and nutritional characteristics (Fig. 4). The PCA plot showed distinct clustering between high and low CH_4_ producing accessions, with the first two components (Dim1 and Dim2) together explaining 87.6% of the total variation, with 77.4% and 10.2% of the variation captured by Dim1 and Dim2, respectively. High CH_4_-yielding accessions were grouped on the negative side of Dim1 and positively associated with higher values of CH_4_ (CH_4_ yield and gas volume), VFA (acetate, propionate, butyrate and valerate), CP and digestibility (IVOMD)-related traits. In contrast, low CH_4_-yielding accessions were located on the positive side of Dim1 and linked with greater values of structural carbohydrates (NDF and ADF) as well as the acetate: propionate ratio. These variables were negatively associated with CH_4_-production traits. These findings show that differences in CH_4_ production among accessions were closely associated with variation in nutrient composition and rumen fermentation characteristics.

**Fig. 4.**
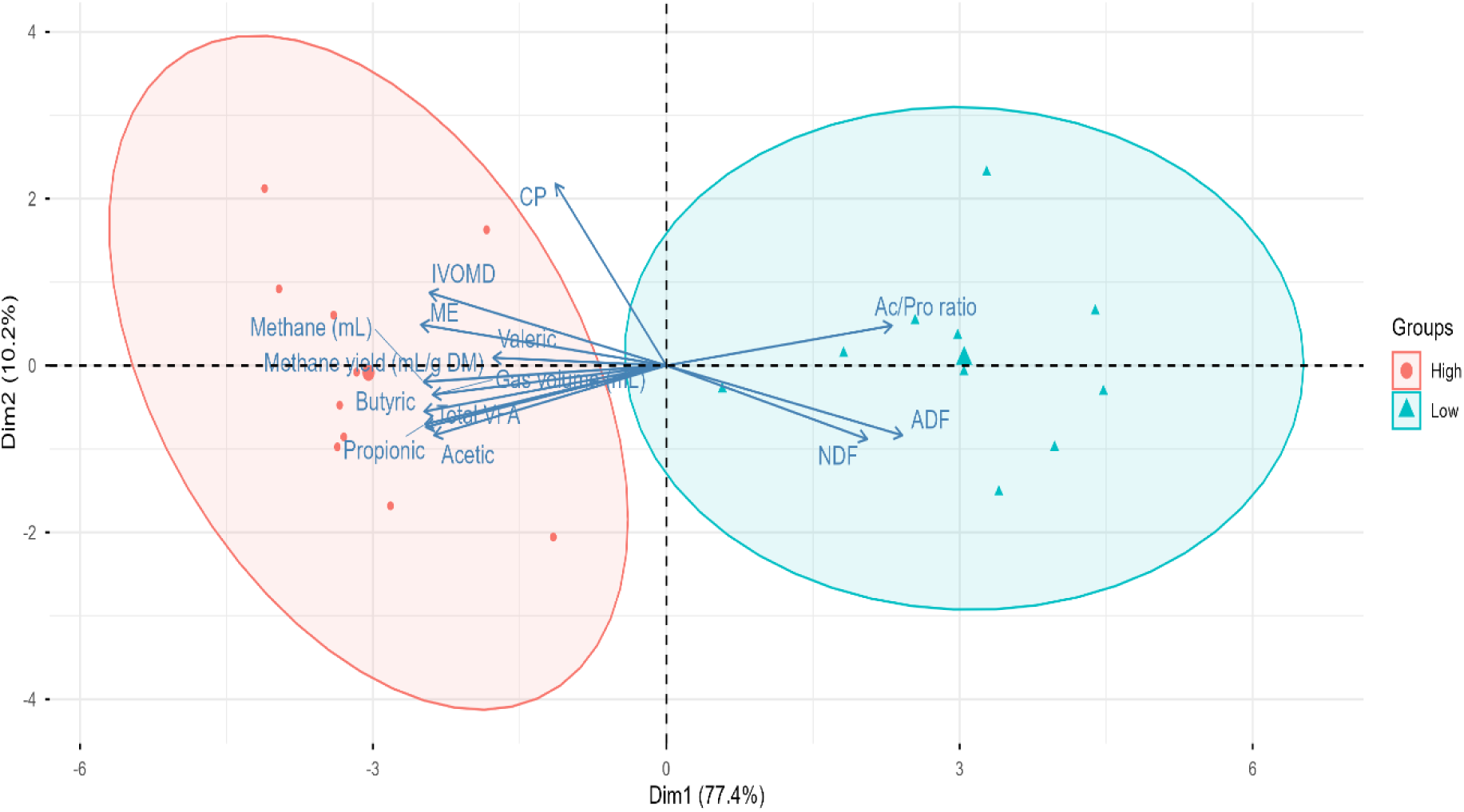
Principal component analysis (PCA) of NG accessions based on CH_4_ yield, and associated nutritional and fermentation traits, showing clustering of high and low CH_4_ yielding accessions. High = accessions with higher CH_4_ yield, Low = accessions with lower CH_4_ yield, CP: crude protein, NDF: neutral detergent fibre, ADF: acid detergent fibre, ME: metabolizable energy, IVOMD: in vitro organic matter digestibility, Ac/Pro = acetate: propionate ratio.

### 3.5. Validation of CH_4_ phenotypes across rumen fluid donors

Accessions exhibiting extreme phenotypes (the top 10 [high] and bottom 10 [low] CH_4_ producers) selected solely based on CH_4_ yield were further examined for consistency in CH_4_ production across three biological replicates using a 120-mL Wheaton bottle assay (as described above). In donor 1, CH_4_ yield among low-and high-CH_4_ producing accessions ranged from 6.1 to 19.8 mL/g DM (mean = 11.6 mL/g DM) and from 21.5 to 41.2 mL/g DM (mean = 33.1 mL/g DM), respectively (Fig. 5). Similarly, in donor 2, low-and high-CH_4_ yielding accessions varied from 5.9 to 16.3 mL/ g DM (mean = 11.4 mL/g DM) and from 19.1 to 33.8 mL/g DM (mean = 26.7 mL/g DM), respectively (Fig. 5). In donor 3, CH_4_ yield ranged from 4.6 to 23.2 mL/g DM (mean = 11.2 mL/g DM) for low-CH_4_ yielding accessions and from 24.5 to 43.9 mL/g DM (mean = 33.6 mL/g DM) for high-CH_4_ producing accessions, respectively (Fig. 5). The ranking of accession was highly consistent across biological replicates, with low-CH_4_ yielding accessions consistently maintaining lower CH_4_ production, whereas high-CH_4_ yielding accessions remained among the highest CH_4_ producers across all donors, with the correlation coefficients of 0.88, 0.91 and 0.86 between screening and validation CH_4_ yield across donors 1, 2 and 3, respectively (p < 0.001, Fig. 5). Among the low CH_4_ producing accessions, accession 16816 consistently produced the lowest CH_4_ yield across all biological replicates, resulting in reductions of 71%, 69% and 79% compared with the corresponding animal means in donors 1, 2, and 3, respectively.

**Fig. 5.**
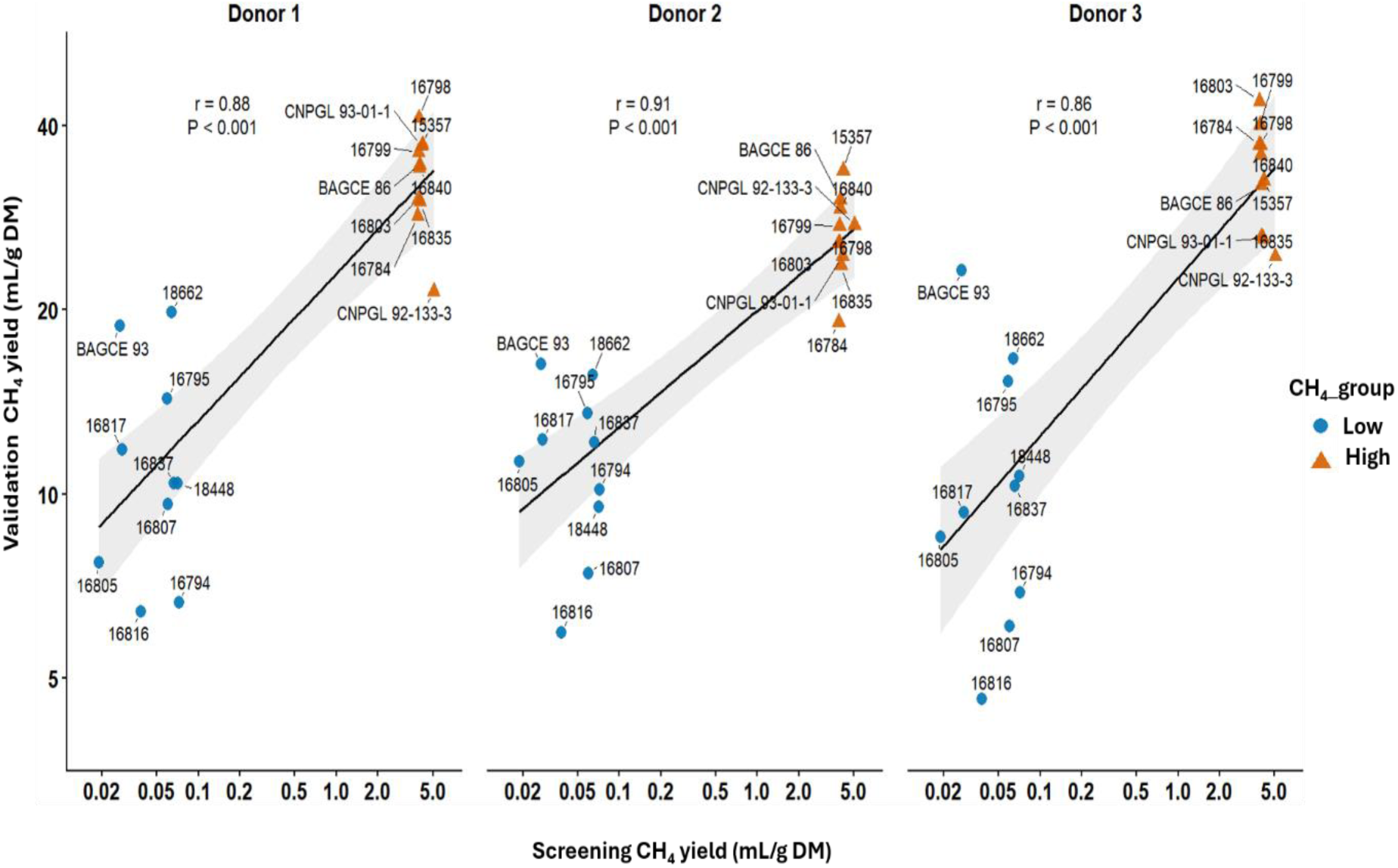
Relationships between screening CH_4_ yield (mL/g DM) and validation CH_4_ yield (mL/g DM) of NG accessions using independent rumen inoculum from three donor animals. High = accessions with higher CH_4_ yield, Low = accessions with lower CH_4_ yield. Screening CH_4_ yield represents the initial CH_4_ yield measured in the high-throughput 20-mL vial assay with pooled rumen fluid, which was used to classify accessions as low-or high-CH_4_ emitters. Validation CH_4_ yield refers to the CH_4_ yield subsequently measured for the selected low-and high-CH_4_ accessions using a larger volume 120-mL Wheaton bottle assay with rumen inoculum obtained independently from three donor animals. Each point represents an individual accession, with accession names shown next to the corresponding points. Regression lines represent linear relationships, with shaded areas indicating 95% confidence interval.

### 3.6. Identification of elite Napier grass accessions using a multi-trait selection index

In addition to selecting accessions with extreme phenotypes solely based on CH_4_ yield (as presented above), a multi-trait selection combining both CH_4_ yield and IVOMD was further employed to identify accessions with relatively low CH_4_ production while maintaining acceptable digestibility levels (≥ 55%). Results of the multi-trait selection index identified ten elite (low-CH_4_ yielding/high IVOMD) and ten poor-performing (high-CH_4_ yielding/low IVOMD) accessions (Table 3). Elite accessions produced lower CH_4_ yield (0.019–0.810 mL/g DM) and had relatively high digestibility (55.2–69.0% IVOMD). The weighted selection index ranked accession 16621 as the best overall due to its low CH_4_ yield (0.384 mL/g DM) coupled with high IVOMD (66.1%). Some other elite accessions with relatively high rankings included 16834, CNPGL 92-198-7, 16838 and 1026 which all had IVOMD ≥ 67% and CH_4_ yields < 0.90 mL/g DM. Interestingly, although several accessions initially categorised as low-CH_4_ emitters based exclusively on CH_4_ yield did not remain low emitters when digestibility was incorporated, accessions 16805, 16795, 18448 and BAGCE 93 remained consistently low after applying a multi-trait selection that combine both CH_4_ yield and IVOMD.

**Table 3.**
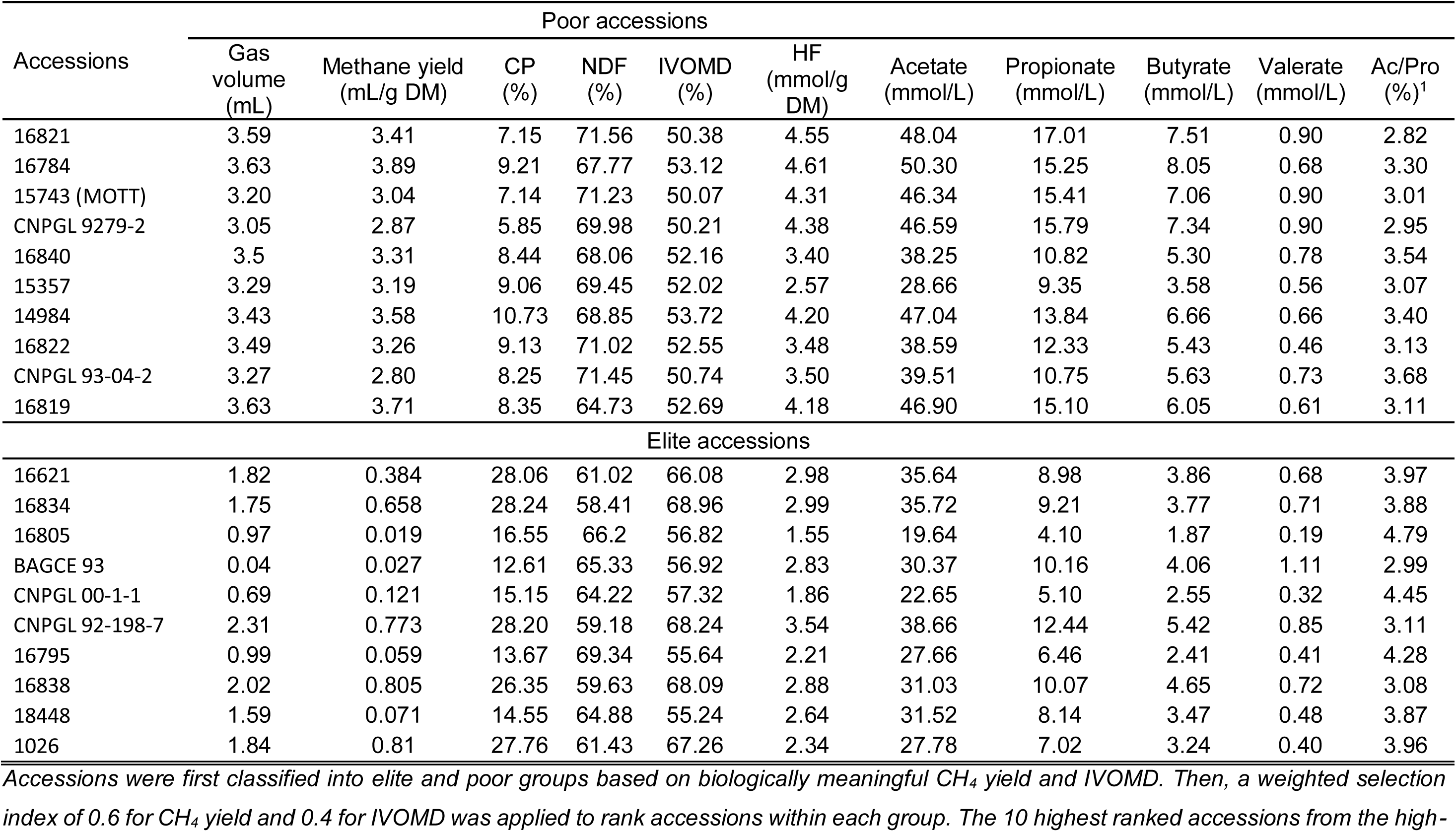
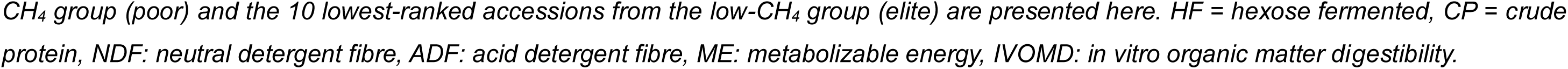
Top and bottom 10 NG accessions ranked using a multi-trait selection under contrasting seasonal and water stress conditions.

| Accessions | Poor accessions |  |  |  |  |  |  |  |  |  |  |
| --- | --- | --- | --- | --- | --- | --- | --- | --- | --- | --- | --- |
|  | Gas volume (mL) | Methane yield (mL/g DM) | CP (%) | NDF (%) | IVOMD (%) | HF (mmol/g DM) | Acetate (mmol/L) | Propionate (mmol/L) | Butyrate (mmol/L) | Valerate (mmol/L) | Ac/Pro (%)<sup>1</sup> |
| 16821 | 3.59 | 3.41 | 7.15 | 71.56 | 50.38 | 4.55 | 48.04 | 17.01 | 7.51 | 0.90 | 2.82 |
| 16784 | 3.63 | 3.89 | 9.21 | 67.77 | 53.12 | 4.61 | 50.30 | 15.25 | 8.05 | 0.68 | 3.30 |
| 15743 (MOTT) | 3.20 | 3.04 | 7.14 | 71.23 | 50.07 | 4.31 | 46.34 | 15.41 | 7.06 | 0.90 | 3.01 |
| CNPGL 9279-2 | 3.05 | 2.87 | 5.85 | 69.98 | 50.21 | 4.38 | 46.59 | 15.79 | 7.34 | 0.90 | 2.95 |
| 16840 | 3.5 | 3.31 | 8.44 | 68.06 | 52.16 | 3.40 | 38.25 | 10.82 | 5.30 | 0.78 | 3.54 |
| 15357 | 3.29 | 3.19 | 9.06 | 69.45 | 52.02 | 2.57 | 28.66 | 9.35 | 3.58 | 0.56 | 3.07 |
| 14984 | 3.43 | 3.58 | 10.73 | 68.85 | 53.72 | 4.20 | 47.04 | 13.84 | 6.66 | 0.66 | 3.40 |
| 16822 | 3.49 | 3.26 | 9.13 | 71.02 | 52.55 | 3.48 | 38.59 | 12.33 | 5.43 | 0.46 | 3.13 |
| CNPGL 93-04-2 | 3.27 | 2.80 | 8.25 | 71.45 | 50.74 | 3.50 | 39.51 | 10.75 | 5.63 | 0.73 | 3.68 |
| 16819 | 3.63 | 3.71 | 8.35 | 64.73 | 52.69 | 4.18 | 46.90 | 15.10 | 6.05 | 0.61 | 3.11 |
|  | Elite accessions |  |  |  |  |  |  |  |  |  |  |
|  | Gas volume (mL) | Methane yield (mL/g DM) | CP (%) | NDF (%) | IVOMD (%) | HF (mmol/g DM) | Acetate (mmol/L) | Propionate (mmol/L) | Butyrate (mmol/L) | Valerate (mmol/L) | Ac/Pro (%)<sup>1</sup> |
| 16621 | 1.82 | 0.384 | 28.06 | 61.02 | 66.08 | 2.98 | 35.64 | 8.98 | 3.86 | 0.68 | 3.97 |
| 16834 | 1.75 | 0.658 | 28.24 | 58.41 | 68.96 | 2.99 | 35.72 | 9.21 | 3.77 | 0.71 | 3.88 |
| 16805 | 0.97 | 0.019 | 16.55 | 66.2 | 56.82 | 1.55 | 19.64 | 4.10 | 1.87 | 0.19 | 4.79 |
| BAGCE 93 | 0.04 | 0.027 | 12.61 | 65.33 | 56.92 | 2.83 | 30.37 | 10.16 | 4.06 | 1.11 | 2.99 |
| CNPGL 00-1-1 | 0.69 | 0.121 | 15.15 | 64.22 | 57.32 | 1.86 | 22.65 | 5.10 | 2.55 | 0.32 | 4.45 |
| CNPGL 92-198-7 | 2.31 | 0.773 | 28.20 | 59.18 | 68.24 | 3.54 | 38.66 | 12.44 | 5.42 | 0.85 | 3.11 |
| 16795 | 0.99 | 0.059 | 13.67 | 69.34 | 55.64 | 2.21 | 27.66 | 6.46 | 2.41 | 0.41 | 4.28 |
| 16838 | 2.02 | 0.805 | 26.35 | 59.63 | 68.09 | 2.88 | 31.03 | 10.07 | 4.65 | 0.72 | 3.08 |
| 18448 | 1.59 | 0.071 | 14.55 | 64.88 | 55.24 | 2.64 | 31.52 | 8.14 | 3.47 | 0.48 | 3.87 |
| 1026 | 1.84 | 0.81 | 27.76 | 61.43 | 67.26 | 2.34 | 27.78 | 7.02 | 3.24 | 0.40 | 3.96 |

Accessions were first classified into elite and poor groups based on biologically meaningful CH<sub>4</sub> yield and IVOMD. Then, a weighted selection index of 0.6 for CH<sub>4</sub> yield and 0.4 for IVOMD was applied to rank accessions within each group. The 10 highest ranked accessions from the high-
*CH<sub>4</sub> group (poor) and the 10 lowest-ranked accessions from the low-CH<sub>4</sub> group (elite) are presented here. HF = hexose fermented, CP = crude* *protein, NDF: neutral detergent fibre, ADF: acid detergent fibre, ME: metabolizable energy, IVOMD: in vitro organic matter digestibility.*

In contrast, poor-performing accessions had higher CH_4_ yields (2.80–3.89 mL/g DM) and lower digestibility (50.1–54.7% IVOMD). Accessions with the least favourable combination of traits were 16821 and 16784 with high CH_4_ yields (>3.40 mL/g DM) and low IVOMD values (<54%). Comparison of chemical composition and fermentation characteristics revealed that elite accessions had lower NDF (58.4–69.3%) concentrations and higher CP (12.6–28.2%) concentrations than poor-performing accessions, which had low CP concentrations (5.9– 10.7%). Metabolisable energy was also higher in elite accessions compared with poor-performing accessions (7.5–9.3 and 7.0–7.7 MJ/kg DM, respectively). Similarly, elite accessions also had lower total VFA production (26.5–50.7 mmol/L) as well as lower concentrations of acetate, propionate and butyrate than poor-performing accessions (42.6– 75.6 mmol/L). The nutritional composition and fermentation characteristics were also significantly different between the two groups (Table 4). For instance, elite accessions had greater CP (p < 0.001), ME (p < 0.001) and IVOMD (p < 0.001), and lower NDF (p = 0.001), CH_4_ yield (p < 0.001), acetate (p = 0.002), propionate (p = 0.001) and butyrate (p = 0.001) concentrations, than poor-performing accessions.

**Table 4.** Mean nutritional compositional differences between top and bottom CH_4_ groups identified using a multi-trait selection.

| Parameters | Poor accessions | Elite accessions<sup>1</sup> | SEM<sup>2</sup> | P-value |
| --- | --- | --- | --- | --- |
| CP (%) | 8.33 | 21.11 | 1.607 | <0.001 |
| NDF (%) | 69.41 | 62.96 | 0.931 | 0.001 |
| ADF (%) | 44.73 | 33.87 | 1.405 | <0.001 |
| ME (MJ/ kg DM) | 7.26 | 8.42 | 0.181 | <0.001 |
| IVOMD (%) | 51.77 | 62.06 | 1.385 | <0.001 |
| Gas volume (mL) | 3.41 | 1.40 | 0.163 | <0.001 |
| Methane yield (mL/g DM) | 3.31 | 0.37 | 0.112 | <0.001 |
| Hexose fermented (mmol/g DM) | 3.92 | 2.58 | 0.198 | 0.002 |
| Methane intensity (mmol/mmol) | 0.04 | 0.01 | 0.002 | <0.001 |
| Acetate (mmol/L) | 43.02 | 30.07 | 1.985 | 0.002 |
| Propionate (mmol/L) | 13.56 | 8.17 | 0.810 | 0.001 |
| Butyrate (mmol/L) | 6.26 | 3.53 | 0.382 | 0.001 |
| Valerate (mmol/L) | 0.72 | 0.59 | 0.071 | 0.212 |
| Ac/Pro (%)<sup>2</sup> | 3.20 | 3.84 | 0.149 | 0.021 |
| Total VFA (mmol/L) | 64.60 | 43.47 | 3.206 | 0.002 |

<sup>1</sup>Accessions were first classified into poor and elite groups based on biologically meaningful CH<sub>4</sub> yield and IVOMD. Then, a weighted selection index of 0.6 for CH<sub>4</sub> yield and 0.4 for IVOMD was applied to rank accessions within each group. The 10 highest ranked accessions from the high-CH<sub>4</sub> group (poor) and the 10 lowest-ranked accessions from the low-CH<sub>4</sub> group (elite) were included in the analysis. <sup>2</sup>Values are expressed as standard error of the mean, <sup>2</sup>acetic:propionate ratio, Abbreviations: CP: crude protein, NDF: neutral detergent fibre, ADF: acid detergent fibre, ME: metabolizable energy, IVOMD: in vitro organic matter digestibility.

### 3.7. Estimated hexose fermentation and methane intensity

Estimated HF varied substantially among accessions, ranging from 1.37 to 5.67 mmol/g DM, with a mean value of 3.14 mmol/g DM. There was significant difference between low-and high-CH_4_ accessions in HF (p < 0.001). Low-CH_4_ accessions exhibited lower HF (mean = 2.16 mmol/g DM) compared with high-CH_4_ accessions (mean = 3.86 mmol/g DM). Among the low-CH_4_ producing accessions, accession BAGCE 93 had higher HF (2.8 mmol/g DM), followed by accessions 18448 (2.6 mmol/g DM) and 16794 (2.4 mmol/g DM), whereas accessions 16784 (4.6 mmol/g DM) and 16798 (4.4 mmol/g DM) showed greatest HF among high-CH_4_ producing accessions.

Methane produced per unit of fermented hexose (CH_4_ intensity) differed among NG accessions, varying from 0.001 to 0.09 mmol CH_4_/mmol HF. High CH_4_-yielding accessions ranged from 0.04 to 0.09 mmol CH4/mmol HF, whereas low-CH_4_ producing accessions ranged approximately from 0.001 to 0.003 mmol CH4/mmol HF. As a result, CH_4_ production per mmol of fermented carbohydrate was consistently lower in low-CH_4_ accessions compared to high-CH_4_ accessions. Among the low-CH_4_ accessions, 16805 had the lowest CH_4_ intensity, followed by 16816 and 16817. Accessions with the highest CH_4_ intensities belonged to the high CH_4_ producing groups, include CNPGL 92-133-3 and 16784.

Methane yield was positively associated with estimated HF (r = 0.55, p < 0.001, Fig. 6). As carbohydrate fermentation increased, CH_4_ production generally increased as well. While some low-CH_4_ accessions exhibited some degree of carbohydrate fermentation, the CH_4_ generated remained at relatively low levels.

**Fig. 6.**
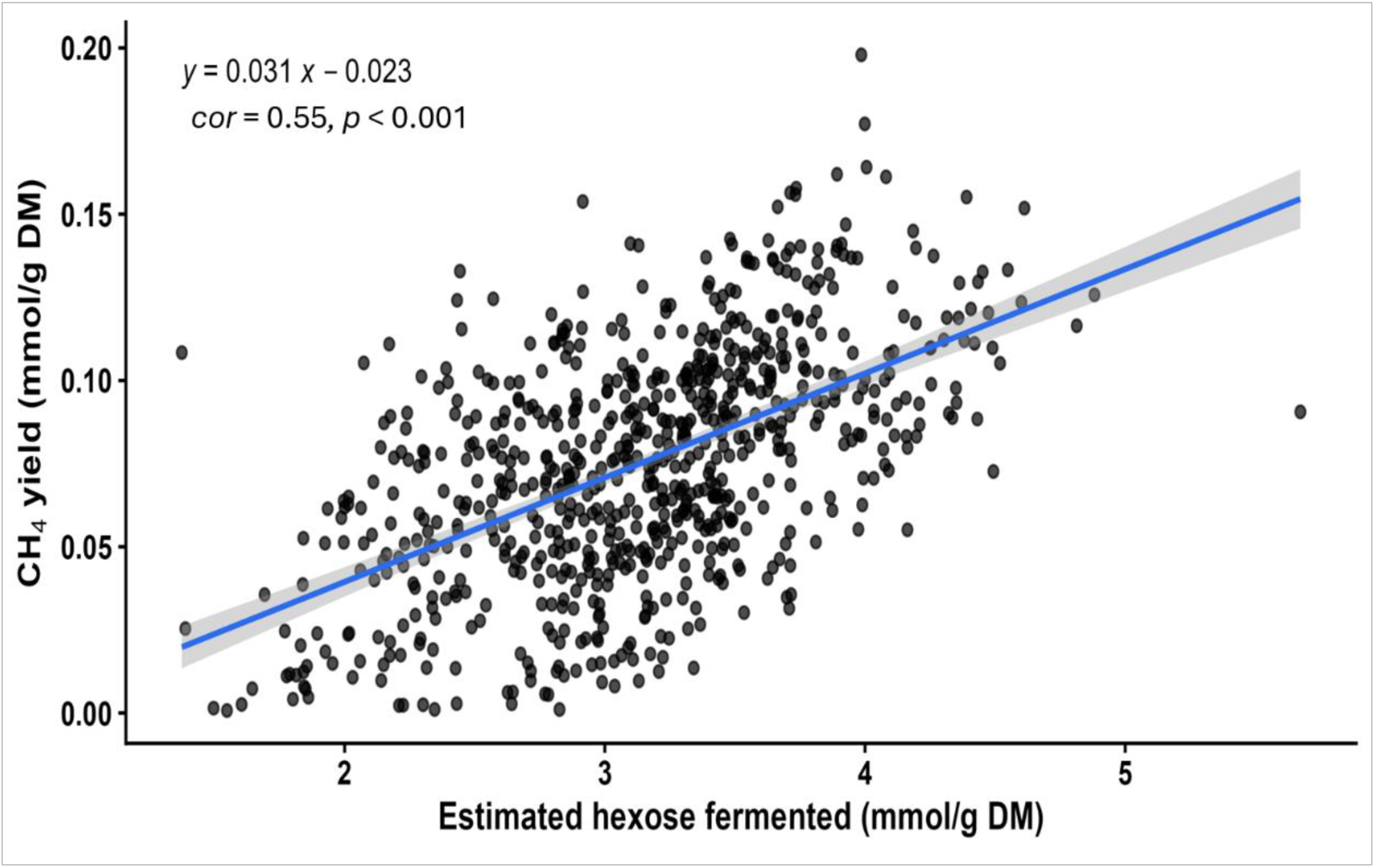
Relationships between CH_4_ yield and hexose fermentation in the incubation medium. Methane production showed a positive association with hexose fermentation, indicating that greater amounts of fermented hexose were accompanied by increased CH_4_ production.

### 3.8. Stability and transitions of methane production among NG accessions across growing conditions

In this case, accessions were divided into three CH_4_-emission groups (low, medium and high) based on tertile-based thresholds (as described above). The alluvial plot demonstrates that there was large variation in CH_4_-emission ranks between growing conditions (Fig. 7). While several accessions shifted between low, medium and high groups across growing conditions, some accessions remained consistent within the same CH_4_-emission category across environments. Accessions that remained consistently low across multiple growing conditions had stable rankings as low emitters, while stable high-emitting accessions generally stayed high across conditions (Supplementary Table 5). For instance, accessions 16621, 16838, 18448, 18438, 16782, 16902, BAGCE 93, and BAGCE 34 exhibited consistently low-emission across growing conditions, whereas accessions 16806, 15357, CNPGL 92-133-3, BAGCE 86, 16815, 16803, 16821, and CNPGL 93-37-5 were consistently high-emitters across growing conditions. Even though a large fraction of accessions changed classification between environments, especially between dry vs. wet season, CH_4_ yield was generally lower in the wet season under SWS.

**Fig. 7.**
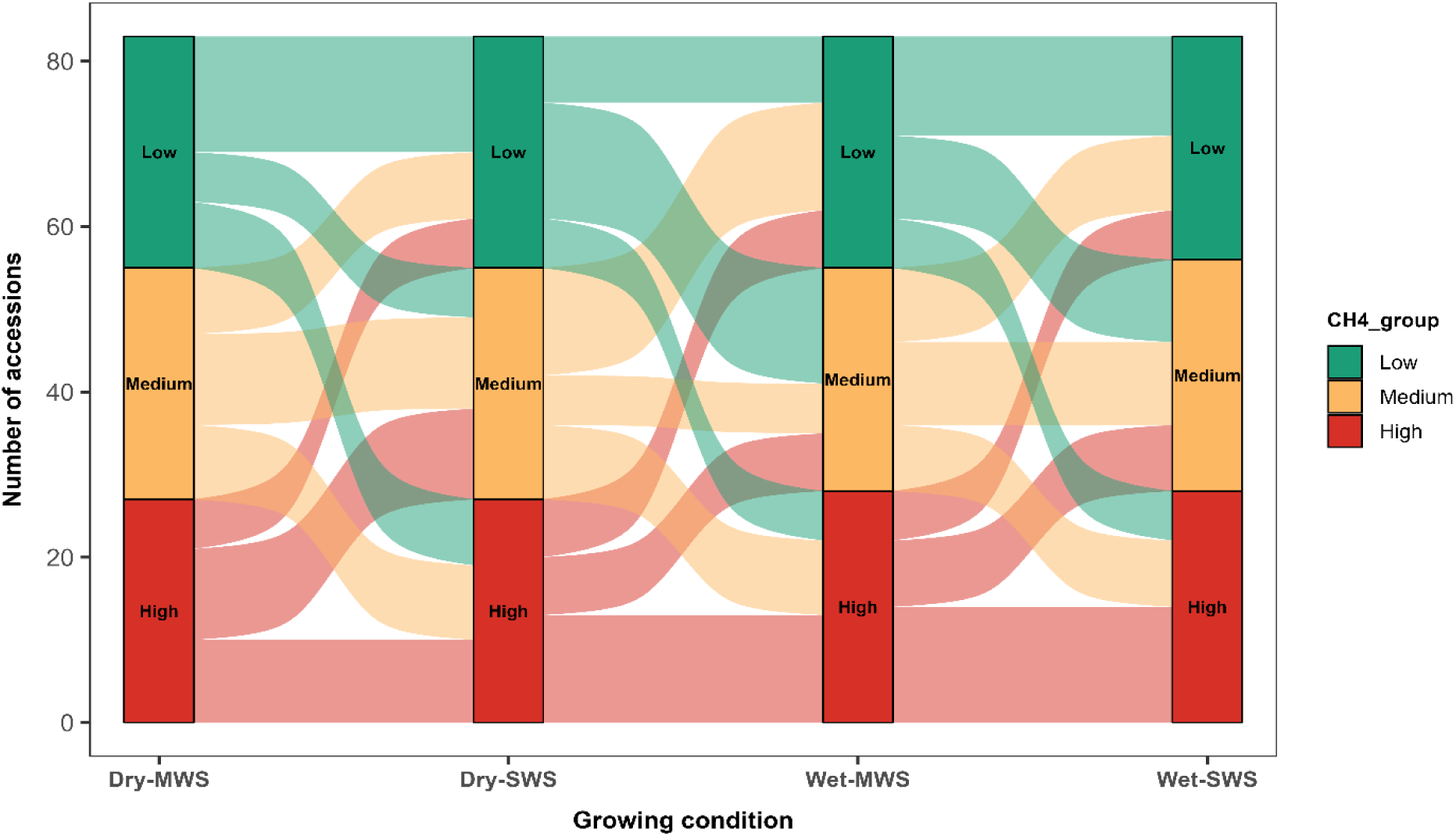
Alluvial plot illustrating transitions in CH_4_-emission groups of NG accessions across four growing conditions (Dry-MWS, Dry-SWS, Wet-MWS, and Wet-SWS). Within each growing condition, accessions were ranked according to CH_4_ yield and classified into low, medium, or high groups using tertile-based thresholds. Flows represent individual accessions and depict changes in CH_4_-emission classification between growing conditions. Oscillating flows indicate fluctuations in CH_4_-emission groups between environments, whereas uninterrupted flows within the same group indicate stable CH_4_ production across growing conditions. MWS = medium water stress; SWS= severe water stress; Dry and Wet refer to the growing season.

## 4. Discussion

This study used high-throughput *in vitro* fermentation to characterise CH_4_ production from a large collection of NG accessions grown under contrasting seasonal and water-stress conditions. Methane yield varied substantially among accessions and environments, and the ranking of extreme phenotypes was broadly retained when tested with inoculum from individual donor cows. Season affected CH_4_ production more strongly than water stress, while multi-trait selection identified accessions combining low CH_4_ yield with acceptable digestibility. These findings establish a practical first-stage phenotyping framework for incorporating CH_4_-related traits into NG improvement.

### 4.1. Large phenotypic variation among NG accessions provides opportunities for methane mitigation

Methane yield ranged from 0.02 to 5.07 mL/g DM, a greater than 250-fold difference, and the overall coefficient of variation was 47%. This wide distribution demonstrates substantial phenotypic variation in methanogenic potential within the NG collection. Previous studies have reported variation among forage species or small numbers of cultivars in digestibility, fibre composition, fermentation and CH_4_ production (Archimède et al., 2011; Pellikaan et al., 2011; Navarro-Villa et al., 2013; Lee et al., 2017; Loza et al., 2021; Jayasinghe et al., 2022). The present study extends this evidence through large-scale, within-species screening and identifies candidate accessions for further evaluation.

The significant accession effect after accounting for season, water stress and block indicates that differences in CH_4_ production were not exclusively attributable to the growing environment. This result is consistent with accession-specific variation in the chemical and physical properties of the substrate entering the rumen. Forage improvement offers a means of altering the substrate available for rumen fermentation and could complement selection of low-CH_4_ animals, dietary supplementation and microbiome-based approaches (Roehe et al., 2016; de Haas et al., 2021; Beauchemin et al., 2022).

Several accessions retained similar CH_4_ rankings across contrasting growing conditions, whereas others moved between low-, medium-and high-producing groups. The former provides candidates for stable performance; the latter show that relative rankings can change even though the mixed model did not detect significant season × accession or water-stress × accession interactions. Environment-dependent rankings are common for forage yield and quality because climate and soil conditions affect growth, cell-wall development and nutritive composition, with subsequent effects on rumen fermentation (Buxton and Fales, 1994; Casler, 1999). Stable favourable phenotypes are particularly valuable where cultivars will be deployed across variable tropical environments (Annicchiarico, 2002; Yan and Kang, 2003).

The multi-trait approach addressed the fundamental limitation of selection based on CH_4_ yield alone: poorly fermentable, highly fibrous accessions may appear favourable simply because less substrate is degraded. Combining CH_4_ production with IVOMD identified accessions with low methanogenic potential, acceptable digestibility and moderate fibre concentrations. This partial separation of CH_4_ yield from feeding value is consistent with evidence linking CH_4_ production to forage fibre and fermentation patterns while retaining exploitable variation among forages (Pinares-Patiño et al., 2007; Beauchemin et al., 2008; Hristov et al., 2013).

### 4.2. Environmental conditions influenced methane production

Season had a stronger effect on CH_4_ production than water stress. Across water-stress treatments, CH_4_ yield was higher in the dry season than in the wet season. Dry-season forage nevertheless had lower mean NDF and ADF concentrations and higher IVOMD than wet-season forage. Fibre concentration alone therefore did not explain the seasonal response. The wet-season material also produced less total VFA, acetate, propionate and butyrate, indicating a lower overall extent of fermentation during the fixed 24-h incubation. Positive associations between CH_4_ yield and these fermentation end-products support the interpretation that seasonal differences in substrate degradation, rather than a simple fibre-CH_4_ relationship, were important under the conditions tested (Moss et al., 2000; Janssen, 2010).

Seasonal changes in temperature, radiation, rainfall and soil moisture can alter plant maturity, leaf-to-stem ratio, cell-wall development and water soluble-carbohydrate concentration, thereby changing digestibility and rumen fermentation (Hsu et al., 1985; Buxton and Fales, 1994; Wilson, 1994; Van Soest, 1994; Archimède et al., 2011; Beauchemin et al., 2022). Greater cell-wall deposition during slow growth or stress may reduce microbial access to fermentable polysaccharides (Jung and Allen, 1995). Environmental effects on phenolics, flavonoids and tannins could also contribute because these compounds can inhibit methanogens, protozoa or broader microbial activity (Patra and Saxena, 2009; Jayanegara et al., 2012; Selmar and Kleinwächter, 2013).

Water stress had no significant main effect on CH_4_ yield, despite its known effects on forage yield and nutritive value (Habte et al., 2022). Responses of forage quality to drought vary with genotype, developmental stage, and the intensity and duration of stress (Dumont et al., 2015; Volaire, 2018). Moderate deficits can delay maturity and cell-wall deposition, whereas prolonged or severe stress can accelerate senescence and lignification (Buxton and Fales, 1994; Jung and Allen, 1995). The absence of an overall response may therefore reflect contrasting effects among accessions or seasons, or the relative water-stress tolerance of NG. The significant season × water-stress interaction confirms that water restriction cannot be interpreted independently of the wider growing environment. Together with changes in accession ranking, this result supports multi-environment evaluation to identify both stable and specifically adapted germplasm (Casler, 1999; Annicchiarico, 2002).

### 4.3. Relationships between forage nutritional composition, digestibility and methane production

Correlation analysis and PCA linked variation in CH_4_ production to forage composition, digestibility and fermentation. Methane yield was positively associated with IVOMD, ME, total gas and total VFA, but negatively associated with NDF and ADF. These associations suggest that the amount of substrate degraded during incubation was a major determinant of CH_4_ yield. More digestible material supports greater microbial fermentation, VFA formation and production of H_2_ and CO_2_ for methanogenesis (Hungate, 1966; Moss et al., 2000; Janssen, 2010). Comparable relationships have been reported for both tropical and temperate forages (Pinares-Patiño et al., 2003; Archimède et al., 2011; Beauchemin et al., 2008).

The accessions selected solely for high CH_4_ yield also had greater IVOMD, ME, gas production and VFA concentrations than the low-CH_4_ group. Greater access to structural and non-structural carbohydrate can enhance microbial growth and fermentation but also increases the supply of reducing equivalents to methanogens (Van Soest, 1994; Moss et al., 2000; McAllister and Newbold, 2008; Pereira et al., 2022). This trade-off is central to forage-based mitigation: a reduction in absolute CH_4_ production is not beneficial if it results only from reduced digestion and lower energy availability to the animal. Alternative H_2_ sinks, or plant traits that alter fermentation pathways without constraining digestibility, are therefore required to lower CH_4_ intensity rather than fermentation itself.

The negative associations between CH_4_ yield and NDF or ADF are consistent with slower fermentation of structural carbohydrate enclosed within lignified cell walls (Van Soest, 1994; Jung and Allen, 1995; Hatfield and Fukushima, 2005). At a fixed 24-h endpoint, accessions with more fibre would be expected to produce less VFA and CH_4_ because less substrate had been degraded. Similar inverse associations have been reported for perennial ryegrass, maize silage and tropical grasses (Getachew et al., 2004; Archimède et al., 2011). Lower CH_4_ production from these accessions should not be interpreted as an inherently favourable phenotype: they also had lower IVOMD, ME and gas production, which could reduce intake and productive performance (Van Soest, 1994; McDonald et al., 2011, van Gastelen et al., 2026).

Positive associations of CH_4_ yield with total VFA, acetate, propionate and butyrate further indicate that CH_4_ production largely reflected the overall extent of fermentation. The VFAs are major products of carbohydrate degradation and supply approximately 70% of metabolisable energy to ruminants (Bergman, 1990); their production is accompanied by reducing equivalents that support methanogenesis (Moss et al., 2000; Hook et al., 2010; Janssen, 2010). Both acetate and propionate production were higher in high-CH_4_ accessions, and the proportional increase in propionate was greater, reducing the acetate-to-propionate ratio. Thus, absolute VFA production was more informative than the acetate-to-propionate ratio when comparing these diverse substrates (Morvay et al., 2011; Janssen, 2010). Importantly, some accessions combined low CH_4_ yield with acceptable IVOMD and moderate fibre, providing evidence that the fermentation-quality trade-off can be reduced through multi-trait selection.

### 4.4. Methane intensity and hexose fermentation reveal differences in fermentation efficiency

The extent of carbohydrate fermentation is a primary driver of CH_4_ production, but the amount of CH_4_ formed per unit of fermented substrate is also relevant when screening germplasm. Estimated HF was positively associated with CH_4_ yield, as expected from the production of VFA, CO_2_ and reducing equivalents during carbohydrate fermentation (Demeyer, 1991; Moss et al., 2000). Methane intensity also differed among accessions. Similar relationships between CH_4_ production and fermentation characteristics have been reported for forage and mixed diets (Ellis et al., 2007; Archimède et al., 2011; Ramin and Huhtanen, 2013).

Methane intensity ranged from 0.001 to 0.09 mmol CH_4_/mmol HF, indicating that variation in CH_4_ yield was not explained entirely by estimated fermentation. Several low-CH_4_ accessions produced less CH_4_ per unit of HF than high-CH_4_ accessions, which is consistent with less reducing equivalents reaching methanogenesis. In the rumen, H_2_ may also be used in propionate formation, microbial biomass synthesis, reductive acetogenesis and other H_2_-consuming pathways (Janssen, 2010; Ungerfeld, 2015, 2020). Differences in cell-wall accessibility, carbohydrate supply and secondary metabolites could affect microbial colonisation and electron flow, while interactions between methanogens and competing H_2_ users could further modify CH_4_ formation (Van Soest, 1994; Greening et al., 2019; Vasta et al., 2019; Ungerfeld, 2020).

### 4.5. Multi-trait selection improves identification of low-CH_4_ emitting Napier grass genotypes

Selection on CH_4_ yield alone identified accessions with low methanogenic potential but also risked retaining material with low digestibility and feeding value. Incorporating IVOMD into the weighted index allowed mitigation and nutritional value to be considered simultaneously. Most elite accessions had substantially lower CH_4_ yield while retaining acceptable to high IVOMD, showing that reduced CH_4_ production was not invariably coupled to poor digestibility. The composition of elite and poor-performing accessions illustrates their contrasting phenotypes. Elite accessions contained more than twice as much CP, less NDF and ADF, mean IVOMD was 62.1% compared with 51.8% in the poor-performing group. Higher CP commonly reflects a greater leaf proportion and can support microbial growth, whereas greater cell-wall concentration restricts microbial attachment and fibre degradation (Van Soest, 1994; Jung and Allen, 1995; Minson, 2012). Relationships between lower fibre, improved digestibility and greater energy availability have also been reported for tropical grasses, including NG (Lee et al., 2017; Negawo et al., 2017).

Despite their greater IVOMD, elite accessions produced less total gas and total VFA than poor-performing accessions, with corresponding reductions in acetate, propionate and butyrate. Methane declined proportionally more than VFA, and elite accessions produced less CH_4_ per unit of estimated HF. These patterns are consistent with differences in both the extent of fermentation and the apparent partitioning of fermentation products (Janssen, 2010; Ramin and Huhtanen, 2013; Ungerfeld, 2020). The higher acetate-to-propionate ratio in the elite group does not contradict this interpretation because concentrations of both acids were lower and the ratio alone does not represent total electron flow.

### 4.6. Validation across donor animals supports repeatable methane phenotypes

High-throughput screening is useful only if selected phenotypes are repeatable with independent inocula. Retesting of the highest-and lowest-CH_4_ yielding accessions using rumen fluid from three donor cows changed the absolute magnitude of fermentation but broadly retained the contrast between groups; accession 16816 was consistently among the lowest producers. High-CH_4_ accessions also produced more total gas, total VFA, acetate, propionate and butyrate across donors. The repeatable ranking supports a substrate-driven component to the observed phenotype and argues against its being an artefact of a single pooled inoculum. Comparable retention of substrate rankings despite variation in absolute output has been reported in other *in vitro* studies (Mould et al., 2005; van Gelder et al., 2005; Serment et al., 2016; Yáñez-Ruiz et al., 2016).

Variation among donors was expected because rumen communities differ in taxonomic composition and fermentative capacity even among animals receiving the same diet (Henderson et al., 2015; Tapio et al., 2017). Inoculum source can therefore alter absolute gas, VFA and CH_4_ measurements (Mould et al., 2005; Ramin et al., 2015; Yáñez-Ruiz et al., 2016). The stability of the group contrast indicates that forage composition and digestibility exerted a consistent influence across these microbial backgrounds (Van Soest, 1994; McAllister et al., 1994).

Agreement between the rapid 20-mL vial assay and the larger common deployed Wheaton-bottle system further supports the use of high-throughput phenotyping to rank large sample sets. The principal cleaner-production advantage is resource efficiency at the discovery stage: the assay reduces analytical labour, incubation space and material requirements before costly field and animal evaluation (Mauricio et al., 1999; Rymer et al., 2005). Accessions retaining their phenotype in the validation assay can now be prioritised for replicated breeding trials and mechanistic assessment.

### 4.7. Implications for climate-smart forage production

Climate smart forage production seeks to prevent environmental burdens while improving the efficiency with which resources are used. Low-CH_4_ forage cultivars could act upstream in the livestock production chain by embedding mitigation in the feed resource rather than relying solely on recurrent supplementation. This may be particularly relevant in tropical and subtropical systems, where forage dominates ruminant diets and animals are often difficult to supplement consistently (Rao et al., 2015; Paul et al., 2020). The present results identify a route towards that objective, but CH_4_-production potential alone is not evidence of lower whole-system impacts. Biomass yield, land and water requirements, animal productivity and life-cycle greenhouse-gas emissions must also be evaluated.

The wide accession-level variation and identification of accessions combining low CH_4_ yield with acceptable digestibility support an integrated breeding strategy. Methane-related traits should be evaluated alongside biomass yield, forage quality, persistence, drought tolerance and disease resistance rather than treated as stand-alone objectives (Casler and Van Santen, 2010; Hayes et al., 2013; Caradus and Chapman, 2025). The strong seasonal effect and changes in relative accession rankings show that selection from a single growing condition could be misleading, despite the absence of significant season × accession and water-stress × accession interactions. Candidate germplasm must therefore be tested across representative sites and seasons, with stability incorporated explicitly into the selection index. Accessions that maintain low CH_4_ production and high nutritional value across environments are the strongest candidates for developing broadly adapted cultivars; responsive accessions may still be useful for specific production zones.

Forage breeding could complement established mitigation options. Feed additives such as 3-nitrooxypropanol and *Asparagopsis* spp. can substantially reduce enteric CH_4_, but regular delivery, cost and availability may constrain use in grazing and resource-limited systems (Vyas et al., 2018; Kinley et al., 2020; Roque et al., 2021; Beauchemin et al., 2022). Once developed and adopted, low-CH_4_ cultivars would not require daily dosing and could reach dispersed livestock populations through seed or planting-material systems.

## 5. Conclusions

This study demonstrated that there is a substantial phenotypic variation for CH_4_ production among NG accessions, thus enabling the identification of forage cultivars with contrasting CH_4_-emission potential. The broad range of CH_4_ yield and associated variation in forage composition and fermentation characteristics showed distinct differences in CH_4_ production among NG germplasms. Additionally, variation observed across different growing conditions highlighted the importance of evaluating candidate genotypes under contrasting environments rather than relying on measurements obtained from a single growing condition. The observation that several accessions maintained consistently low CH_4_ phenotypes across seasons and water-stress conditions further suggests that accession-specific differences could be stable and a useful basis for future breeding and germplasm evaluation.

The independent validation provided further support for the initial CH_4_-based screening. Accessions classified as low-and-high emitters solely on the basis of CH_4_ yield were subsequently evaluated using rumen inoculum from individual donor cows. Despite differences in CH_4_ production among donor animals, the contrast between the low-and high-CH_4_ accessions was generally maintained, indicating that the observed differences were not limited to a single rumen inoculum source. These results support the use of CH_4_ production as a potential selection trait, although confirmation under animal feeding trial is still needed.

The subsequent multi-trait selection addressed a limitation of selecting accessions based on CH_4_ production alone: a low-CH_4_ phenotype is not necessarily sufficient if it is accompanied by a poor forage nutritive value. Combining CH_4_ yield with IVOMD therefore allowed the identification of elite accessions that combine low CH_4_ yield with relatively high digestibility, providing useful candidates for further forage evaluation compared with selection based solely on CH_4_ production. Their contrasting fermentation profiles, including differences in CH_4_ production relative to fermented substrate, also suggest that variation in CH_4_ output may be related to differences in fermentation efficiency and not simply to the amount of substrate fermented. In summary, this stepwise approach, combining large-scale germplasm screening, validation and multi-trait selection, identify a subset of NG germplasm with potential value for developing lower-CH_4_ forage cultivars without compromising forage digestibility and provides a practical basis for considering CH_4_-related traits in future forage improvement programmes and in the development of more sustainable livestock production systems.

## Author contributions

**Agalu W. Zeleke**: conceptualisation, methodology, investigation, laboratory analysis, data processing and curation, statistical analysis, visualisation, and original manuscript writing. **Juan Palma-Hidalgo**: Investigation, laboratory analysis, manuscript review and editing. **Zaira Pardo Dominguez:** Investigation, laboratory analysis, manuscript review and editing. **Meki Muktar:** data collection, manuscript review and editing. **Abel Teshome**: data collection, manuscript review and editing. **Bayissa Hatew**: data collection, manuscript review and editing. **Chris S. Jones:** fund acquisition, project administration, manuscript review and editing. **C. Jamie Newbold:** conceptualisation, methodology, investigation, fund acquisition, project administration, manuscript review and editing. All authors have read and approved the final manuscript.

## Acknowledgements

This study is funded as part of the UK-CGIAR Centre. The UK-CGIAR Centre aims to support global food security by bringing together scientists from the UK and the CGIAR to form impact-focused research collaborations. The Centre is funded by UK International Development from the UK government. The authors also acknowledge EMBRAPA for making its germplasm and breeding lines available for this study.

## Data availability

All data generated and analysed during this study are available upon request.

