## Supplementary materials for "Large-scale phenotyping and multi-trait selection identify low-methane Napier grass (*Cenchrus purpureus*) accessions under contrasting environmental conditions"

Supplementary Table 1. Mean and range of feed quality traits and rumen fermentation characteristics of NG accessions under different growing conditions

| Traits | Dry-MWS | Mean | Dry-SWS | Mean | Wet-MWS | Mean | Wet-SWS | Mean |
| --- | --- | --- | --- | --- | --- | --- | --- | --- |
| CP (%) | 4.30–20.92 | 9.79±0.19 | 7.99–28.24 | 19.02±0.36 | 5.84–19.82 | 9.57±0.16 | 7.09–18.17 | 11.45±0.15 |
| NDF (%) | 60.61–69.56 | 64.63±0.13 | 57.27–67.63 | 62.69±0.14 | 59.32–73.85 | 68.27±0.21 | 61.43–74.60 | 68.19±0.17 |
| ADF (%) | 29.6–43.34 | 38.08±0.15 | 25.27–40.29 | 32.11±0.24 | 37.37–49.70 | 44.37±0.15 | 36.63–48.75 | 42.64±0.16 |
| ADL (%) | 2.27–3.77 | 3.03±0.02 | 1.98–3.71 | 2.94±0.02 | 2.06–4.24 | 3.14±0.03 | 2.08–4.21 | 3.08±0.03 |
| IVOMD (%) | 50.73–64.88 | 54.94±0.16 | 53.65–68.96 | 62.37±0.24 | 48.85–59.03 | 52.70±0.14 | 49.22–57.66 | 53.72±0.11 |
| ME (MJ/kg DM) | 7.07–8.92 | 7.67±0.02 | 7.49–9.37 | 8.61±0.03 | 6.76–8.04 | 7.30±0.02 | 6.78–7.87 | 7.40±0.01 |
| OM (%) | 76.64–85.81 | 81.29±0.12 | 77.85–86.34 | 83.11±0.01 | 77.47–86.31 | 82.68±0.11 | 79.04–87.49 | 83.37±0.09 |
| Gas volume (mL) | 0.04–3.83 | 2.82±0.04 | 1.52–4.03 | 2.87±0.04 | 1.49–3.59 | 2.60±0.03 | 0.32–3.63 | 2.15±0.05 |
| CH_4_ yield (mL/g DM) | 0.03–4.04 | 2.20±0.05 | 0.38–5.07 | 2.41±0.07 | 0.39–3.41 | 1.81±0.05 | 0.02–3.89 | 1.30±0.07 |
| Acetate (mmol/L) | 24.21–54.66 | 36.56±0.41 | 22.22–64.79 | 36.45±0.47 | 19.77–48.04 | 34.64±0.41 | 14.05–52.87 | 34.93±0.56 |
| Propionate (mmol/L) | 0.06–15.63 | 10.59±0.15 | 5.47–17.63 | 11.10±0.18 | 5.34–17.01 | 10.28±0.17 | 4.09–15.88 | 10.01±0.20 |
| Butyrate (mmol/L) | 0.15–7.95 | 4.84±0.06 | 2.12–9.21 | 5.05±0.09 | 2.17–7.51 | 4.53±0.08 | 1.61–8.05 | 4.29±0.10 |
| Iso-butyrate (mmol/L) | 0.02–2. 83 | 0.40±0.04 | 0.08–0.77 | 0.47±0.01 | 0.02–0.71 | 0.36±0.02 | 0.02–2.61 | 0.34±0.02 |
| Valerate (mmol/L) | 0.15–1.15 | 0.46±0.03 | 0.24–0.92 | 0.63±0.01 | 0.15–1.02 | 0.51±0.02 | 0.01–1.09 | 0.51±0.02 |
| Iso-valerate (mmol/L) | 0.02–0.85 | 0.35±0.02 | 0.15–1.29 | 0.52±0.01 | 0.03–0.87 | 0.38±0.02 | 0.07–0.86 | 0.42±0.01 |
| Ac/Pro (%) | 2.99–4.23 | 3.45±0.03 | 2.69–4.31 | 3.34±0.03 | 2.77–4.61 | 3.43±0.02 | 2.66–4.79 | 3.57±0.03 |
| Total VFA (mmol/L) | 30.95–79.34 | 53.22±0.58 | 31.60–93.84 | 54.22±073 | 28.37–74.65 | 50.71±0.67 | 23.37–75.95 | 50.49±0.85 |

Abbreviations: neutral detergent fiber (NDF), acid detergent fiber (ADF), acid detergent lignin (ADL), organic matter (OM), crude protein (CP), in vitro organic matter digestibility (IVOMD) and metabolisable energy (ME), CH_4_ = methane

Supplementary Table 2. Significant interaction effects (Season × Accessions) on acetate concentration

| Accession | Estimate (mmol/L) | SE | p-value | Accession | Estimate (mmol/L) | SE | p-value |
| --- | --- | --- | --- | --- | --- | --- | --- |
| 14355 | -14.39 | 6.32 | 0.0233 | 16813 | -16.04 | 6.32 | 0.012 |
| 14389 | -16.92 | 6.63 | 0.011 | 16816 | -23.14 | 6.32 | <0.001 |
| 14982 | -17.77 | 6.32 | 0.005 | 16817 | -14.3 | 6.32 | 0.024 |
| 14983 | -16.92 | 6.32 | 0.007 | 16818 | -13.87 | 5.48 | 0.029 |
| 14984 | -14.34 | 5.48 | 0.009 | 16819 | -20.58 | 5.33 | 0.001 |
| 15357 | -15.82 | 5.48 | 0.004 | 16821 | -13.93 | 5.32 | 0.009 |
| 15743 (MOTT) | -13.07 | 6.32 | 0.039 | 16834 | -18.34 | 6.32 | 0.004 |
| 16621 | -18.15 | 7.07 | 0.011 | 16835 | -15.59 | 5.48 | 0.005 |
| 16782 | -16.83 | 6.32 | 0.008 | 16836 | -16.12 | 6.63 | 0.015 |
| 16783 | -14.13 | 6.32 | 0.026 | 16838 | -17.65 | 6.33 | 0.005 |
| 16784 | -14.76 | 5.48 | 0.007 | 16839 | -15.05 | 6.32 | 0.018 |
| 16785 | -15.67 | 6.32 | 0.014 | 16902 | -15.39 | 6.63 | 0.021 |
| 16786 | -13.12 | 6.32 | 0.039 | 18438 | -16.11 | 6.32 | 0.011 |
| 16788 | -14.76 | 6.32 | 0.02 | BAGCE 17 | -22.41 | 6.32 | <0.001 |
| 16789 | -14.13 | 5.48 | 0.011 | BAGCE 30 | -14.71 | 6.62 | 0.02 |
| 16791 | -12.51 | 5.48 | 0.023 | BAGCE 34 | -19.52 | 6.32 | 0.002 |
| 16792 | -14.01 | 5.48 | 0.011 | BAGCE 81 | -15.84 | 6.3 | 0.013 |
| 16793 | -17.1 | 6.32 | 0.007 | BAGCE 93 | -12.93 | 6.32 | 0.041 |
| 16795 | -16.43 | 5.48 | 0.003 | CNPGL 00-1-1 | -18.63 | 6.23 | 0.003 |
| 16797 | -13.05 | 6.32 | 0.034 | CNPGL 92-198-7 | -17.59 | 6.31 | 0.006 |
| 16798 | -17.93 | 6.32 | 0.005 | CNPGL 92-66-3 | -19.01 | 6.25 | 0.003 |
| 16800 | -14.05 | 6.32 | 0.027 | CNPGL 93-01-1 | -23.49 | 6.63 | <0.001 |
| 16805 | -16.64 | 5.48 | 0.003 | CNPGL 93-18-2 | -14.03 | 6.32 | 0.027 |
| 16807 | -21.32 | 6.32 | 0.001 | CNPGL 93 -37-5 | -15.53 | 6.63 | 0.012 |
| 16808 | -14.06 | 6.32 | 0.027 | CNPGL 94-13-1 | -14.79 | 6.32 | 0.019 |
| 16810 | -16.67 | 6.32 | 0.009 |  |  |  |  |

Supplementary Table 3. Significant interaction effects (Season × Accessions) on propionate concentration

| Accession | Estimate (mmol/L) | SE | p-value | Accession | Estimate (mmol/L) | SE | p-value |
| --- | --- | --- | --- | --- | --- | --- | --- |
| 14389 | -5.83 | 2.47 | 0.019 | 16817 | -5.93 | 2.35 | 0.012 |
| 14355 | -5.94 | 2.35 | 0.012 | 16819 | -7.09 | 2.43 | 0.003 |
| 14982 | -4.86 | 2.35 | 0.039 | 16821 | -4.51 | 1.98 | 0.023 |
| 14983 | -5.39 | 2.45 | 0.022 | 16834 | -5.13 | 2.35 | 0.029 |
| 14984 | -4.98 | 2.04 | 0.015 | 16835 | -4.75 | 2.04 | 0.02 |
| 15357 | -4.58 | 2.03 | 0.018 | 16836 | -5.22 | 2.47 | 0.035 |
| 16782 | -5.17 | 2.35 | 0.029 | 16837 | -4.95 | 2.03 | 0.015 |
| 16783 | -5.27 | 2.36 | 0.027 | 16838 | -6.72 | 2.35 | 0.004 |
| 16784 | -5.42 | 2.03 | 0.008 | 18438 | -5.22 | 2.35 | 0.027 |
| 16785 | -4.88 | 2.35 | 0.039 | BAGCE 17 | -7.9 | 2.35 | <0.001 |
| 16788 | -5.19 | 2.35 | 0.023 | BAGCE 30 | -5.91 | 2.37 | 0.012 |
| 16789 | -5.31 | 2.03 | 0.009 | BAGCE 34 | -6.48 | 2.33 | 0.006 |
| 16791 | -5.01 | 2.04 | 0.014 | BAGCE 81 | -5.98 | 2.35 | 0.011 |
| 16792 | -4.13 | 2.02 | 0.043 | BAGCE 97 | -5.28 | 2.09 | 0.025 |
| 16795 | -5.36 | 2.03 | 0.009 | CNPGL 00-1-1 | -7.21 | 2.35 | 0.002 |
| 16798 | -5.64 | 2.35 | 0.017 | CNPGL 92-198-7 | -6.24 | 2.29 | 0.008 |
| 16805 | -5.47 | 2.03 | 0.007 | CNPGL 92-66-3 | -7.09 | 2.35 | 0.003 |
| 16807 | -8.47 | 2.35 | <0.001 | CNPGL 9279-2 | -5.07 | 2.09 | 0.016 |
| 16808 | -5 | 2.35 | 0.034 | CNPGL 93-01-1 | -7.69 | 2.47 | 0.002 |
| 16810 | -6.23 | 2.35 | 0.008 | CNPGL 93-04-2 | -4.68 | 2.35 | 0.047 |
| 16813 | -5.55 | 2.34 | 0.019 | CNPGL 93-18-2 | -5.92 | 2.31 | 0.012 |
| 16816 | -7.47 | 2.32 | 0.002 | CNPGL 93-18-2 | -5.92 | 2.31 | 0.012 |

Supplementary Table 4. Significant interaction effects (Season × Accessions) on butyrate concentration

| Accession | Estimate (mmol/L) | SE | p-value | Accession | Estimate (mmol/L) | SE | p-value |
| --- | --- | --- | --- | --- | --- | --- | --- |
| 14389 | -2.95 | 1.15 | 0.011 | 16819 | -3.21 | 1.09 | 0.004 |
| 14982 | -2.42 | 1.15 | 0.023 | 16835 | -2.34 | 0.95 | 0.014 |
| 14983 | -2.46 | 1.09 | 0.025 | 16836 | -2.79 | 1.15 | 0.017 |
| 15357 | -2.06 | 0.95 | 0.031 | 16837 | -2.21 | 0.95 | 0.02 |
| 16783 | -2.57 | 1.09 | 0.019 | 16838 | -2.76 | 1.09 | 0.012 |
| 16789 | -2.16 | 0.95 | 0.023 | BAGCE 17 | -3.05 | 1.1 | 0.006 |
| 16791 | -1.89 | 0.95 | 0.046 | BAGCE 34 | -2.83 | 1.09 | 0.01 |
| 16792 | -1.95 | 0.95 | 0.041 | BAGCE 81 | -2.26 | 1.1 | 0.04 |
| 16793 | -2.37 | 1.09 | 0.031 | CNPGL 00-1-1 | -2.48 | 1.09 | 0.024 |
| 16798 | -2.41 | 1.1 | 0.029 | CNPGL 92-198-7 | -2.84 | 1.1 | 0.009 |
| 16805 | -2.01 | 0.95 | 0.034 | CNPGL 92-66-3 | -2.97 | 1.09 | 0.007 |
| 16807 | -3.3 | 1.09 | 0.003 | CNPGL 93-01-1 | -3.02 | 1.15 | 0.009 |
| 16810 | -2.55 | 1.09 | 0.021 | CNPGL 93-18-2 | -2.33 | 1.09 | 0.034 |
| 16816 | -3.24 | 1.1 | 0.003 |  |  |  |  |

Supplementary Table 5. Methane yield of Napier grass accessions consistently classified as low or high emitters across contrasting growing conditions.

| CH_4_ group^1^ | Accession | Growing condition | | | |
| --- | --- | --- | --- | --- | --- |
|  |  | Dry-MWS | Dry-SWS | Wet-MWS | Wet-SWS |
| Consistently low | 16621 | 1.563 | 0.384 | 0.872 | 0.893 |
|  | 16838 | 1.946 | 1.890 | 1.367 | 0.983 |
|  | 18448 | 1.154 | 1.833 | 1.394 | 0.603 |
|  | 18438 | 1.613 | 1.445 | 0.972 | 1.381 |
|  | 16782 | 1.476 | 1.831 | 1.833 | 0.701 |
|  | 16902 | 1.806 | 1.682 | 1.603 | 0.344 |
|  | BAGCE 93 | 0.480 | 1.759 | 1.731 | 1.677 |
| Consistently high | 16806 | 2.406 | 2.725 | 2.641 | 1.852 |
|  | 15357 | 2.733 | 3.155 | 2.223 | 2.001 |
|  | CNPGL 92-133-3 | 2.332 | 3.816 | 2.140 | 1.720 |
|  | BAGCE 86 | 2.988 | 3.112 | 1.831 | 1.655 |
|  | 16815 | 2.414 | 2.536 | 2.199 | 1.855 |
|  | 16803 | 2.073 | 2.835 | 2.107 | 1.984 |
|  | 16821 | 2.120 | 3.220 | 2.452 | 1.482 |
|  | CNPGL 93 -37-5 | 2.214 | 3.504 | 2.192 | 1.497 |

^1^Methane (CH_4_) yield values were ranked and divided into three groups of equal size (tertiles) and accessions in the first third of the CH_4_ yield distribution were considered as ‘low’ CH_4_ emitters, middle third accessions were ‘medium’ emitters, and accessions in the top third were ‘high’ emitters. Accessions with consistently low- and -high CH_4_ values across growing conditions from these CH_4_ categories were presented in this table.

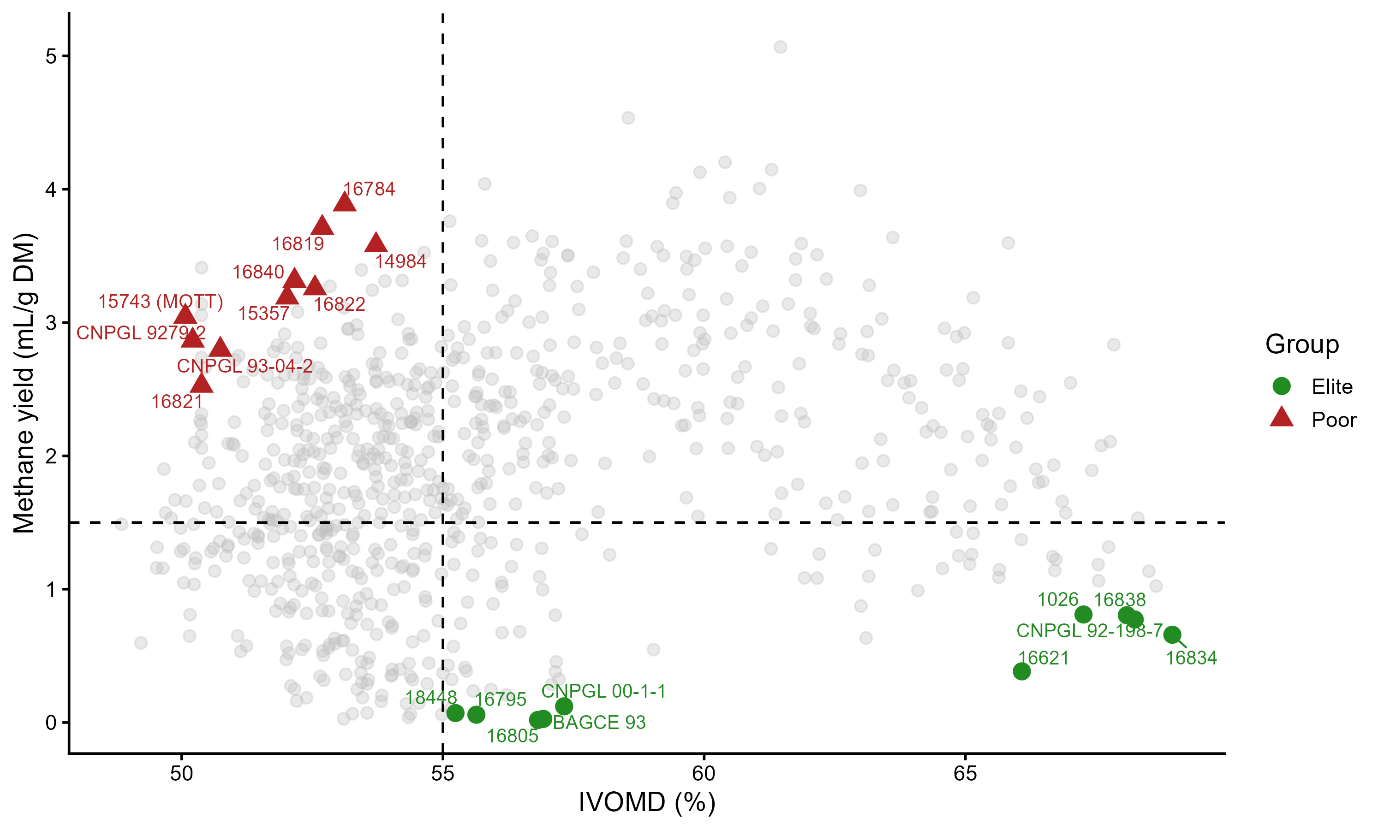

Supplementary Fig. 1. Selection of elite and poor-performing Napier grass accessions based on CH_4_ yield and IVOMD. Scatter plot illustrating the relationship between IVOMD (%) and CH_4_ yield (mL/g DM) for all Napier grass accessions (grey circles). Elite accessions (green circles) were selected based on high IVOMD (>55%) and low CH_4_ yield (≤1.5 mL/g DM), whereas poor accessions (red triangles) exhibited lower digestibility (≤55%) and higher CH_4_ production (>1.5 mL/g DM). Dashed vertical and horizontal lines represent the selection thresholds used to identify contrasting phenotypes for subsequent analyses.
